# Single-cell transcriptomic catalog of mouse cortical development

**DOI:** 10.1101/208744

**Authors:** Lipin Loo, Jeremy M. Simon, Eric S. McCoy, Jesse K. Niehaus, Mark J. Zylka

## Abstract

We generated a single-cell transcriptomic catalog of the developing mouse cerebral cortex that includes numerous classes of neurons, progenitors, and glia, their proliferation, migration, and activation states, and their relatedness within and across timepoints. Cell expression profiles stratified neurological disease-associated genes into distinct subtypes. Complex neurodevelopmental processes can be reconstructed with single-cell transcriptomics data, permitting a deeper understanding of cortical development and the cellular origins of brain diseases.

---

Development of the mammalian cerebral cortex involves a complex series of cell proliferation, differentiation, and migration events. Genetic and environmental factors that perturb any one of these processes can impair intellect and increase risk for neurodevelopmental disorders such as autism spectrum disorder (ASD). Mice are routinely used to study neurodevelopmental processes and to model brain disorders. However, we currently lack a comprehensive understanding of which cells are present in the normally developing mouse cerebral cortex. Here, we transcriptionally profiled a total of 18,545 mouse neocortical cells at two key times of corticogenesis using Drop-seq: 10,931 cells at embryonic day 14.5 (E14.5), representing a progenitor-driven stage, and 7,614 cells at birth (P0), when all six cortical layers are formed and gliogenesis has begun (**Supplementary Fig. 1-2, Supplementary Table 1**)^1,2^. To increase the precision of unbiased clustering, we developed an iterative cell type refinement method (see **Methods**) and identified 22 principal cell types at each age (**Fig. 1a, Supplementary Fig. 3-4, Supplementary Table 2**). Each cell type was classified by expression of known and novel marker genes. We further validated cell type assignment and cortical distribution at each age using *in situ* hybridization data (Eurexpress, Allen Institute of Brain Science, GENSAT) (**Supplementary Fig. 5-14, Supplementary Table 3**). Hierarchical clustering of transcriptomes from each cell type (**Supplementary Table 2**) was used to assess inter-relatedness within and between time points (**Fig. 1a**). We identified distinct cortical layer-specific cell types, which express the longest genes^3^, multiple progenitor-like cell types, including *Eomes*^+^ (*Tbr2*^+^) progenitors, GABAergic interneurons, and non-neuronal cells, such as endothelial cells and microglia. Other cell types from adjacent non-cortical tissues–ganglionic eminences, striatal inhibitory neurons, and thalamus (**Supplementary Fig. 8, 9, 12 and 13**)—were identified and excluded from subsequent analyses.

**Figure 1.**
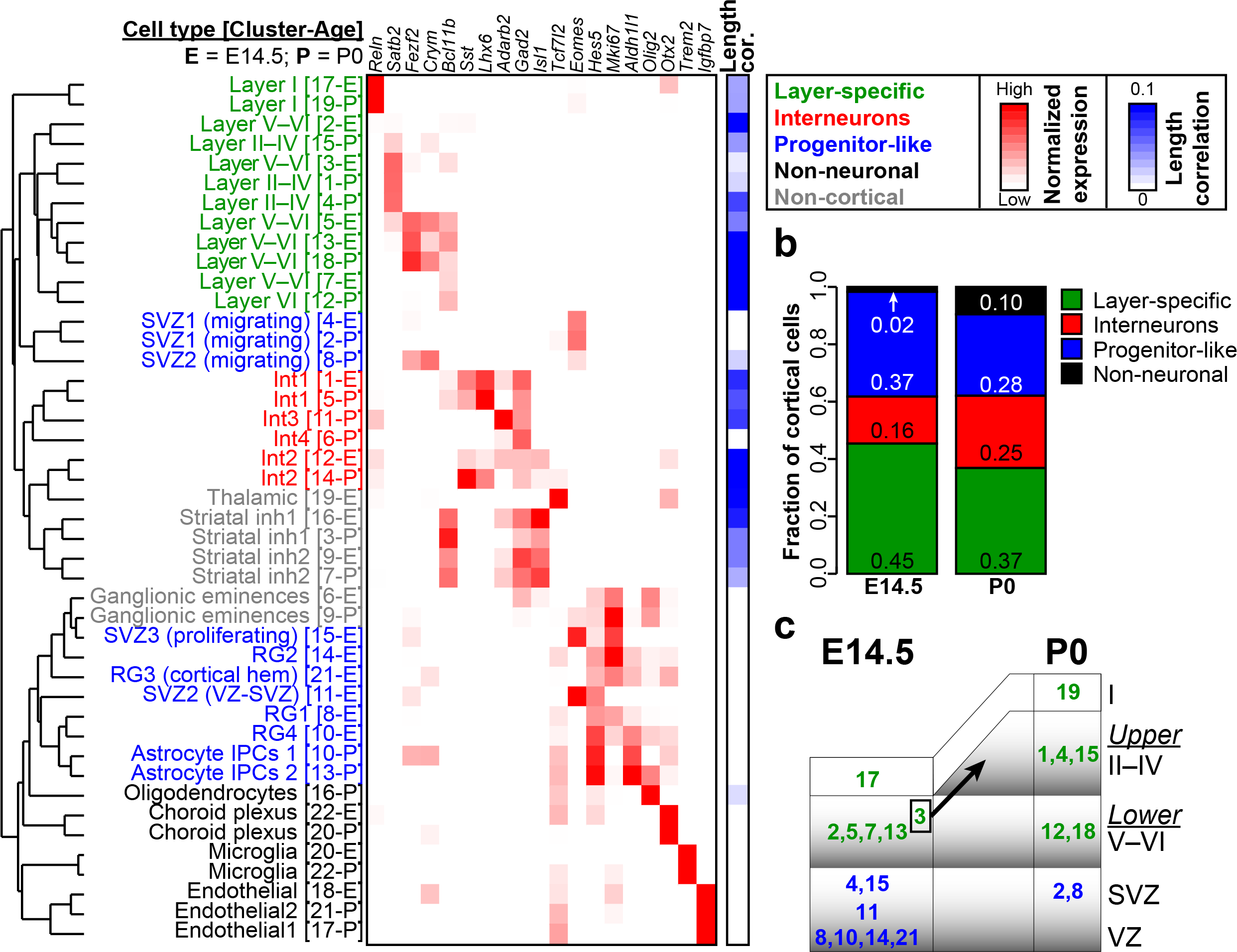
Principal cell types of the developing cerebral cortex identified by single-cell RNA-seq. **a.** Marker genes characterize each cell type in the E14.5 and P0 cerebral cortex. *Mki67* is a marker of cell proliferation. Cell types were grouped into categories (colored), based on their functional identity and transcriptional similarity (Pearson correlation distances, dendrogram). Correlation of expression with gene length provided on a scale of white to blue. **b.** Fractional proportions of cortical cells, averaged across all biological replicates (n=6, E14.5; n=3, P0). Non-cortical cells were excluded. **c.** Cortical layer-specific cells, their spatiotemporal expression and trajectories. Arrow, Cluster 3-E (lower layer neurons; E14.5) is transcriptionally most similar to Cluster 1-P (upper layer neurons; P0).

We grouped the cortical cells into our broad classes (layer-specific, interneurons, progenitor-like, and non-neuronal) and assessed their relative proportions at each age (**Fig. 1b**). Nearly 40% of cells at E14.5 were progenitor-like, including multiple classes of radial glia (RG) and intermediate progenitors localized to the ventricular zone (VZ) and sub-ventricular zone (SVZ), but represented only 28% of cells by P0, as expected^4^. Primates have an additional outer RG layer, marked by *Tnc* and *Ptprz1*. These markers were expressed in mouse progenitor cells (E14.5 RG and P0 Astrocyte IPCs, **Supplementary Fig. 15**) but did not show cell-type specificity, further corroborating that mice lack this distinct layer^5^. Overall, the ratio of excitatory to inhibitory neurons was in line with previous estimates that range from 2:1 to 5:1 (Refs. 6,7). Additionally, the P0 cerebral cortex contained a greater proportion of non-neuronal cells relative to the E14.5 cortex, which is consistent with the known timing of glial proliferation^4^.

Lower-layer neurons were present at E14.5 and were most similar to their P0 counterparts, as expected given the timing of cortical layer formation^2^. One type of lower layer neuron at E14.5 (Layer V-VI; Cluster 3-E) was most similar to an upper layer neuron type at P0 (Layer II-IV; Cluster 1-P) (**Fig. 1a**). This class of lower layer neurons expressed upper (*Satb2*) and lower (*Bcl11b*) layer markers, a migratory marker *Tiam2*, and *Pou3f1*, a transcription factor that is expressed in Layer II-III neurons during their migration and differentiation^2,8^ (**Supplementary Fig. 5, Supplementary Fig. 11**). Together, these data suggest that these cells are destined to become upper-layer neurons (**Fig. 1c**), some of which are born around E14.5 (Ref. 9).

Interneurons migrate tangentially from the ganglionic eminences and populate all layers of the cerebral cortex^10^. Correspondingly, Int1 and Int2 were present at E14.5 and P0 and expressed high levels of *Lhx6*, a transcription factor associated with Parvalbumin (PV) and Somatostatin (SST) interneurons^11^ (**Fig. 1a**). *Sst*, but not *Pv*, was detected in Int1 and Int2 at these ages, as expected^12^. The Int3 and Int4 interneuron classes were found only in the P0 cortex. Int3 (Cluster 11-P) expressed canonical markers of vasoactive intestinal peptide (VIP) interneurons, including *Htr3a*, *Npas1*, and *Adarb2* (Ref. 11). Int4 (Cluster 6-P) expressed *Cdca7*, a marker of some SST^+^ and PV^+^ interneurons^11^, but did not express Sst at this stage. High expression of *Sp9*, *Tiam2*, *Dlx5*—general transcriptional and migratory markers—suggest that Int4 is an immature/migrating interneuron cluster (**Supplementary Fig. 12)**. Together, these data uncover new molecular relationships between cell types and chart the changing cellular landscape during early cortical development.

We next investigated whether within-cluster gene expression heterogeneity could further stratify cell types and cell states. Indeed, by focusing on variable gene expression within clusters, we identified seven sub-clusters at E14.5 and five subclusters at P0 (**Fig. 2**, **Supplementary Fig. 16-17**). These sub-clusters included: 1) cells in various phases of the cell cycle (RG4 [10-E], SVZ3 (proliferating) [15-E] (**Fig. 2b**), ganglionic eminences [6-E, 9-P]), 2) highly related but functionally distinct cell types (e.g. mural, pericyte, and meningeal sub-classes of endothelial cells [18-E, 21-P]), and 3) putative activation states (e.g. among microglia [20-E], oligodendrocytes [16-P], and ganglionic eminences [9-P]). We found no evidence for neuronal activation, as marked by *Fos* and other immediate early genes. Our use of ion channel inhibitors during cell dissociation might thus provide a novel way to block dissociation-induced neuronal immediate early gene induction^13^. We also identified transitioning cell types within Layer I [17-E]: *Fabp7*^+^ precursors, *Ntm*^+^ newborn cells^14^,and *Reln*^+^ mature Cajal-Retzius cells.

**Figure 2.**
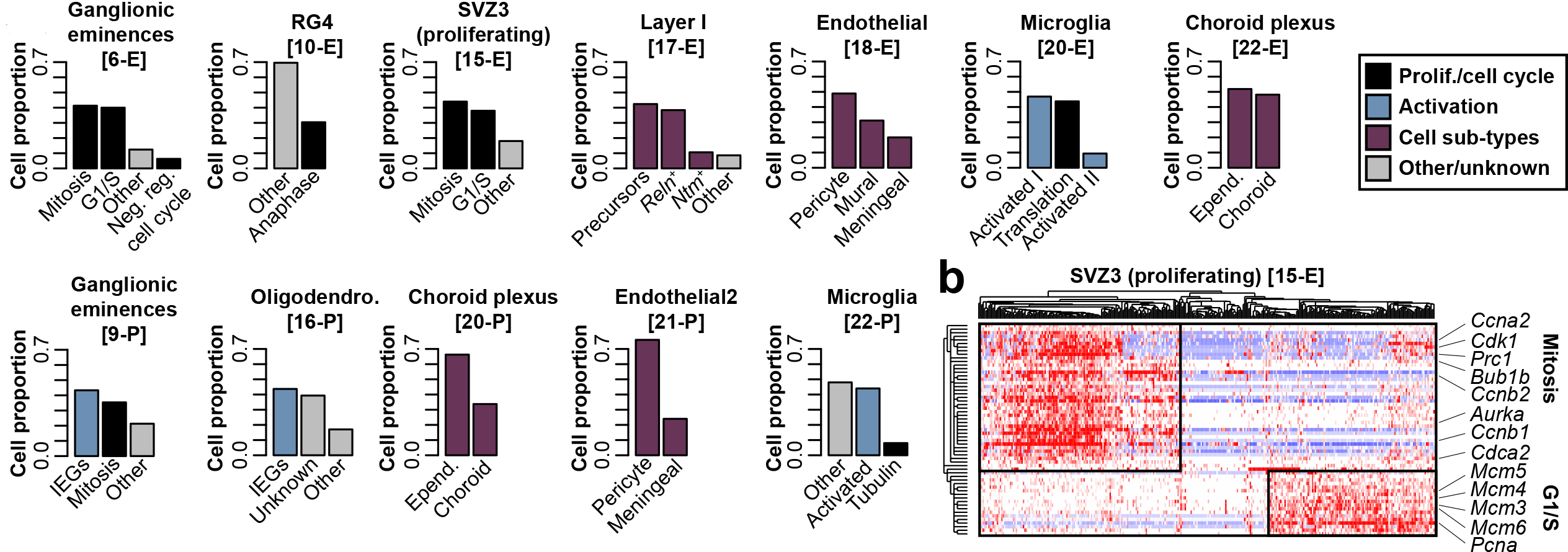
Sub-clustering reveals proliferation and activation states and cell subtypes. **a.** Proportion of cells found within detected sub-clusters at E14.5 (top) and at P0 (bottom). **b.** SVZ3 (cluster 15-E) contains cells in different stages of the cell cycle.

Cortical dysfunction is implicated in neurological and neuropsychiatric diseases, including amyotrophic lateral sclerosis (ALS), Alzheimer’s disease (ALZ), ASD, and schizophrenia (SCZ). Genes that increase risk for these diseases were recently identified^15–18^. To determine if these disease-associated genes are expressed broadly or specifically in developing cortical cell types, we hierarchically clustered the cellular expression profiles of these genes, and defined disease subtypes (**Fig. 3, Supplementary Fig. 18-21,** see **Methods**). The majority of the 14 genes linked to Alzheimer’s disease (e.g. *Apoe*, *Trem2*, *Picalm*, *Cr1l*, *Cd2ap*) were expressed predominantly in non-neuronal cells, especially microglia. Recent studies suggest that microglia contribute to neurodegeneration in Alzheimer’s disease^19^. In contrast, genes linked to other disorders fell into multiple subtypes. For example, ASD Subtype 1 contained numerous synaptic genes, consistent with previous subtyping based on gene ontology and molecular pathway analyses^20^. The remaining five ASD subtypes contained chromatin modifiers and transcriptional regulators, which gene ontology-based methods generally collapse into a smaller number of groups. Thus, cellular expression profiling has the potential to identify novel disease subtypes and cellular vulnerabilities associated with brain diseases.

**Figure 3.**
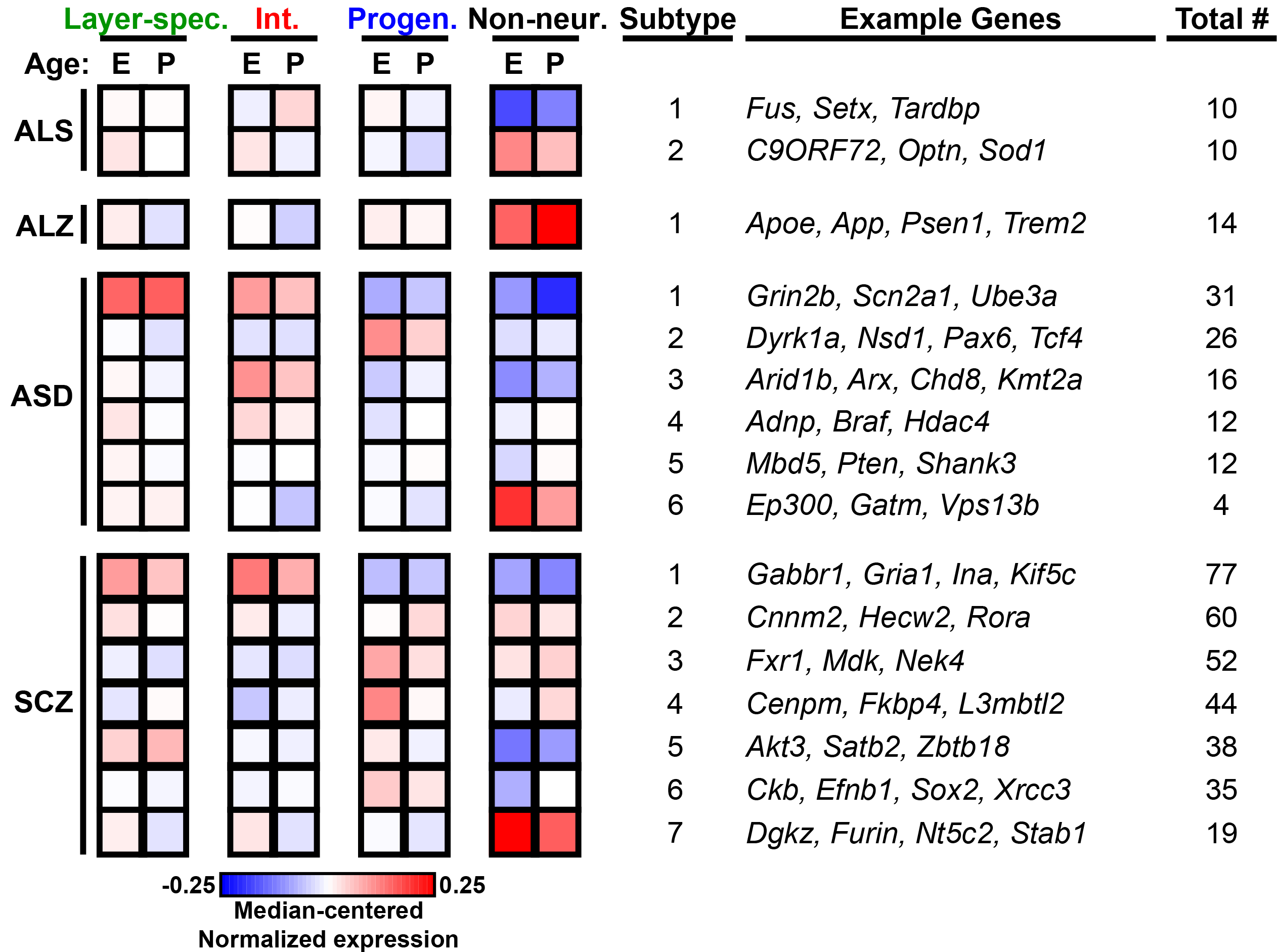
Subtyping neurological diseases with single-cell transcriptomics data. Median-centered gene expression values across all cells within a Cluster and all Clusters within a broad category. Disease-associated genes were hierarchically clustered and subtypes were identified based on distance. Full datasets showing expression of each gene in each cell type is provided in Supplementary Materials.

In conclusion, we found that the unfolding cellular complexity of brain development—formerly pieced together over decades of research^2^—can be reconstructed with a single experimental approach. Additionally, our data reveal novel relationships between cortical cell types over time and cortical space, and provide new insights into the cellular origins and subtypes of neurological diseases. Diverse mouse models show alterations in cortical development, ranging from micro- to macrocephaly to early onset neurodegeneration. Our single-cell data provide an essential resource for future studies directed at understanding how genetic and environmental factors affect cell composition, cell states, and cell fates during early cortical development.

## Acknowledgments

We thank Eva Anton and members of the Zylka lab for helpful discussions and critical reading of the manuscript, Hyejung Won and Jason Stein for discussion of disease subtyping analysis, Marcus Basiri and Garret Stuber for establishing the Drop-seq microfluidics station and discussion of Drop-seq analysis. This research was supported by the National Institute of Environmental Health Sciences of the National Institutes of Health (DP1ES024088, M.J.Z.) and by the Simons Foundation (Award ID # 393316, M.J.Z.). M.J.Z and J.M.S were additionally supported by The Eunice Kennedy Shriver National Institute of Child Health and Human Development (U54HD079124) and National Institute of Neurological Disorders and Stroke (P30NS045892).

## Author Contributions

M.J.Z, L.L, and J.M.S. designed the study. E.S.M. performed cortical dissections. L.L. performed Drop-seq, prepared libraries for high-throughput sequencing and annotation of marker genes. J.M.S. analyzed all Drop-seq data. J.K.N. provided technical assistance. L.L., J.M.S., and M.J.Z. wrote the manuscript.

## Supplementary Materials

### Mouse handling and timed matings

All procedures used in this study were approved by the Institutional Animal Care and Use Committee at the University of North Carolina at Chapel Hill. Mice were maintained on a 12 h:12 h light:dark cycle and given food and water *ad libitum*. Timed matings were set up in the evening, using one male and two female C57Bl/6J mice (Jackson Labs) per breeding cage. The male mouse was separated from female mice the next morning. 14 days after separation, pregnant female mice were euthanized and embryonic day 14.5 (E14.5) embryos were collected for dissections. For P0 pups, cages were monitored daily from E18.5 to postnatal day 0 (P0) for newborn pups.

### Cortical dissections and single cell suspension preparation

Cerebral cortices (both halves) from E14.5 and P0 mice were dissected in neurobasal medium and rinsed with Hank’s Balanced Salt Solution (HBSS; 14175095 Gibco)^1^. Cortices were then incubated for 30 min at 37°C in papain (1 vial diluted with 2.5 mL of HBSS; Pierce 88285) with DNase I (20 mg/mL; D4513 Sigma) in Ca^2+^ and Mg^2+^ free HBSS. Next, 1 mL of neurobasal medium containing 5% FBS was added to the cortical mixture and triturated to deactivate the papain. The cells were centrifuged for 2 min at 4,000 rpm, washed twice, and resuspended in Ca^2+^ and Mg^2+^ free HBSS with ion channel inhibitors (5 μM TTX ab120054 Abcam, 25 μM DL-AP5 ab120004 Abcam, 5 μM DNQX 2312 Tocris). A total of nine replicates were prepared from two developmental time points (6 replicates for E14.5 embryos and 3 replicates for P0 pups). Each replicate contained cortical cells from male and female littermates.

### Drop-seq procedure

Drop-seq was performed largely as described^2^. Briefly, cortical cells were diluted to an estimated concentration of 400 cells/μL in Ca^2+^ and Mg^2+^ free HBSS with ion channel inhibitors and HEK293T cells were spiked in at a concentration of 3% of total cells (12 cells/μL) while barcoded beads (ChemGenes Corporation, catalogue number Macosko201110) were resuspended in lysis buffer to an estimated concentration of 400 beads/μL. Cells and beads were co-encapsulated with oil (Q×200^TM^ Droplet Generation Oil for EvaGreen, Biorad) using a microfluidics chip (Part number 3200455, Dolomite). Droplets of around 3 mL of aqueous volume (1.5 mL of cells and beads) were broken with perfluorooctanol in 30 mL of 6× SSC. The harvested beads were then washed twice with 6x SSC and hybridized RNA was reverse transcribed using Maxima H minus Reverse Transcriptase (ThermoFisher). Populations of 2,500 reverse-transcribed beads (~100 cells) were separately amplified with 13 cycles of PCR (primers, chemistry, and cycle conditions identical to those previously described) and PCR products were purified with 0.6× AMPure XP beads (Agencourt).

cDNA from an estimated 12,000 E14.5 cells and 8,000 P0 cells were pooled, purified and tagmented with Nextera XT DNA Library Preparation kit (Illumina). Input cDNA (1 ng) from each replicate was amplified with custom primer P5_TSO_Hybrid and Nextera index primers (N701, N702, N703, N704, N711, N712, N715, N716 and N718). Tagmented samples were purified twice with 0.6x and 1x AMPure XP beads. All replicates were pooled and sequenced on one Illumina HiSeq 4000 flowcell (eight lanes) to avoid sequencing bias. Read 1 was 20 bp; Read 2 was 50 bp and Read 3 (index) was 8 bp. Data was de-multiplexed using bcl2fastq version 2.18.0.12.

### Processing of Drop-seq data

FASTQ files were converted to BAM format, tagged with cell and molecular barcodes, quality-filtered, trimmed, polyA-trimmed, and converted back to FASTQ as previously described^2,3^. Reads were aligned to a mouse-human hybrid genome (mm10-hg19) using STAR^4^, then sorted, merged, and exon-tagged as described^2,3^. Bead synthesis errors were corrected as described^3^, and BAM files were separated into those containing mouse or human reads. UMIs were determined to be species-specific if >90% of the transcripts came from that species, or considered a doublet if neither species achieved 90% specificity. UMIs were not considered if the transcript count sum (mouse+human) was less than 500. Gene expression matrices were then created using only the mouse-specific UMIs, as described^2,3^. Gene expression matrices were combined from multiple biological replicates; data values for one or more replicates that did not detect a given gene were assigned to zero. Processing steps utilized the Drop-seq Toolkit v1.12 where possible.

### Basic analysis of Drop-seq data

Cells whose mitochondrial contribution exceeded 10% of transcripts were removed, then only genes present in at least 10 cells and having at least 60 transcripts summed across all cells were considered. We then performed batch correction using ComBat^5^ where each independent replicate was considered a batch, thus minimizing any technical variation. To reduce the complexity of the data, we performed Principal Components Analysis (PCA) and eigenvalue permutation (500 shufflings) to determine how many principal components (PCs) to use, as described previously^3^. This yielded 89 PCs for E14.5 and 78 PCs for P0. These data were visualized using t-SNE^6^; we iterated both *perplexity* and *learning*_*rate* parameters to optimize the visualization, ultimately setting these to 50 and 750, respectively. Code made available^3^ was used where possible.

### Cluster identification and refinement

We found that the Louvain-Jaccard clustering method utilized by Shekhar *et al.^3^* produced highly variable results for our data depending on the number of specified nearest neighbors. We therefore devised an iterative procedure that picks the optimal number of nearest neighbors to use for the given dataset. To do this, we iterate from 10 to 100 nearest neighbors, and for each iteration, repeat the Louvain-Jaccard clustering method. Then, we assess how many clusters formed and how robust they were using silhouette widths^7^ based on Spearman correlation distances. After the iteration was complete, we utilized the number of nearest neighbors that produced the maximal average silhouette width across all clusters as a starting point for cell clustering. The silhouette width analysis also allowed us to assess overall performance of each cluster and demonstrated that the basic clustering method published previously^3^ inappropriately assigned many cells to clusters. To refine these cluster assignments, we devised a second iterative approach that attempts to reassign extreme outliers (silhouette width < -0.1) to a better grouping. Over five iterations, these outlier cells of each cluster were given a chance to form their own novel cluster (if its own silhouette width was > 0 and there were at least 10 cells) or join the next-best cluster. If a cell was reassigned to the next-best and remained an outlier there, it would be sent back to its original assignment and flagged such that it would not be considered for reassignment in subsequent iterations. This process improved the overall cluster assignments and resulted in the creation of two novel clusters for E14.5, such that the final number of clusters for both E14.5 and P0 was 22. The final cell type assignments were visualized using t-SNE^6^.

### Identification of cell type marker genes and enriched pathways

Marker genes for each refined cluster were identified as described previously^2,3^, and expression summaries were created using the code provided where possible. To identify biological pathways enriched in each cluster, markers whose expression fold-change relative to other clusters exceeding 0 were mapped to human gene symbols and assessed using a hypergeometric test in Piano^8^ with MSigDB C2 classifications plus additional neurological gene sets as we described previously^9^. Pathways with an FDR < 0.1 and among the top 50 for a given cluster were considered for inclusion in the cell type annotation table.

### ISH validation

Marker genes were validated with *in situ* hybridization data available on Eurexpress (www.eurexpress.org), Allen Brain Institute (www.developingmouse.brain-map.org) and GENSAT (www.gensat.org). Images were cropped to representative sections of the neocortex, ganglionic eminences, striatum, thalamus and choroid plexus. Marker annotations are provided in Supplementary Table 3.

### Cell type sub-clustering method

To identify genes whose expression pattern was heterogeneous within a cluster, we required that a given gene was detected in at least 25% of cells but not more than 75% of cells within a cluster, then further refined the heterogeneous gene list using a feature selection tool^10^ on the expression of all cells within the cluster. The expression of these genes in all cells for that cluster were then clustered hierarchically and inspected manually for coherent patterns representing sub-clusters. Genes within each sub-cluster were then assessed for functional enrichments using ToppGene^11^.

### Cross-age comparisons and disease gene subtyping

Expression data for each cell cluster was merged by taking the mean of all values, and all clusters from both ages were compared to each other using Pearson correlations. To focus on genes most responsible for cell-type-specificity, we used genes that were identified either as cell type markers (log-fold-change > 1.5) or among those that were included in the sub-cluster analysis. Lists of commonly mutated genes for each disease of the cortex were downloaded as follows: ALS (^12^, Tables 1-2), Alzheimer’s disease (^13^, Table 1 + APP, PSEN1, PSEN2), ASD (SFARI Gene classifications Syndromic and 1, obtained August 22, 2017), schizophrenia^14^. For each disease gene set, we performed hierarchical clustering of median-centered expression values across all 44 combined clusters (E14.5 and P0 included) using 1-Pearson correlation distance. We then cut the gene dendrograms at a height of 1.5 to divide genes into subgroups (except ASD, where we instead specified a height of 1.25). Then for each gene subgroup (#genes ≥ 3), we collapsed expression across related cell types by taking the median across those cell types within an age (e.g. median of all E14.5 interneurons) to obtain 8 total values per gene subgroup (layer-specific, interneuron, progenitor-like, and non-neuronal for each age).

**Supplementary Figure 1.**
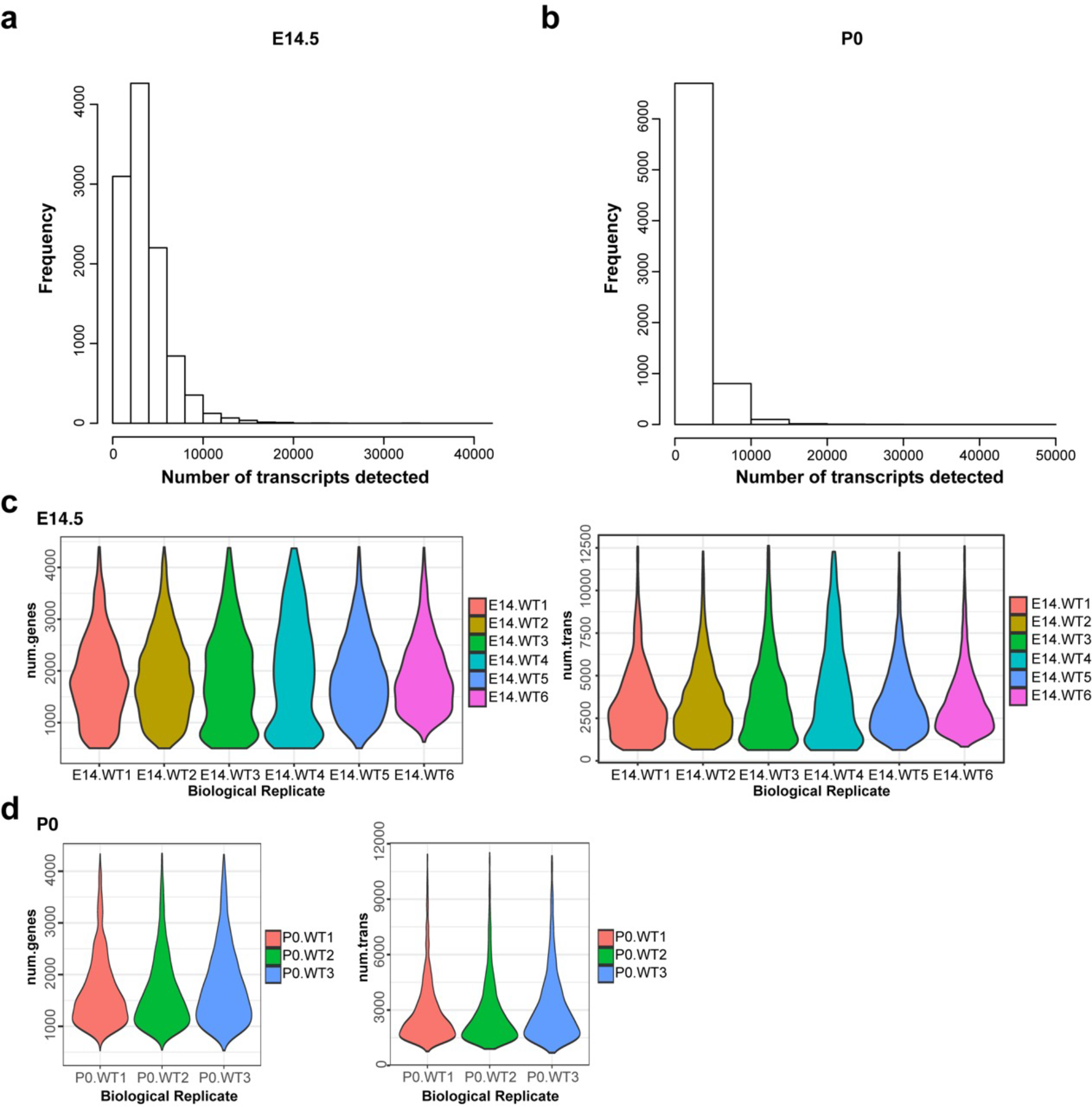
Quality statistics of E14.5 and P0 Drop-seq data. **A-B.** Histogram showing total number of transcripts detected in E14.5 (**a**) and P0 (**b**) samples. **c-d**. Violin plots of total number of genes (left) and total number of transcripts (right) detected for each biological replicate for E14.5 (**c**) and P0 (**d**) samples.

**Supplementary Figure 2.**
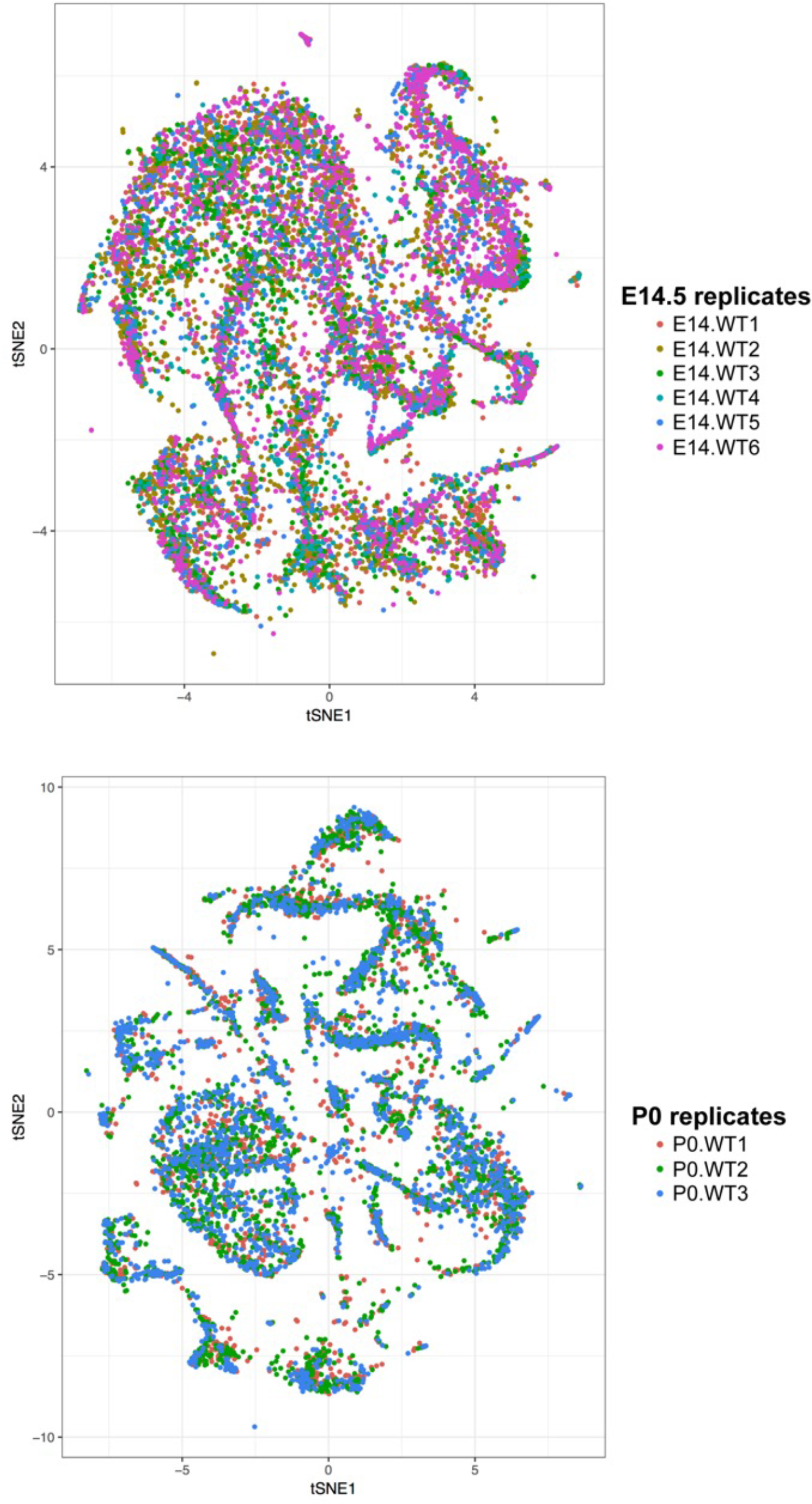
Drop-seq expression data from all replicates visualized by tSNE. tSNE visualization of significant principal components, where points are labeled by their biological replicate for E14.5 (top) and P0 (bottom).

**Supplementary Figure S3.**
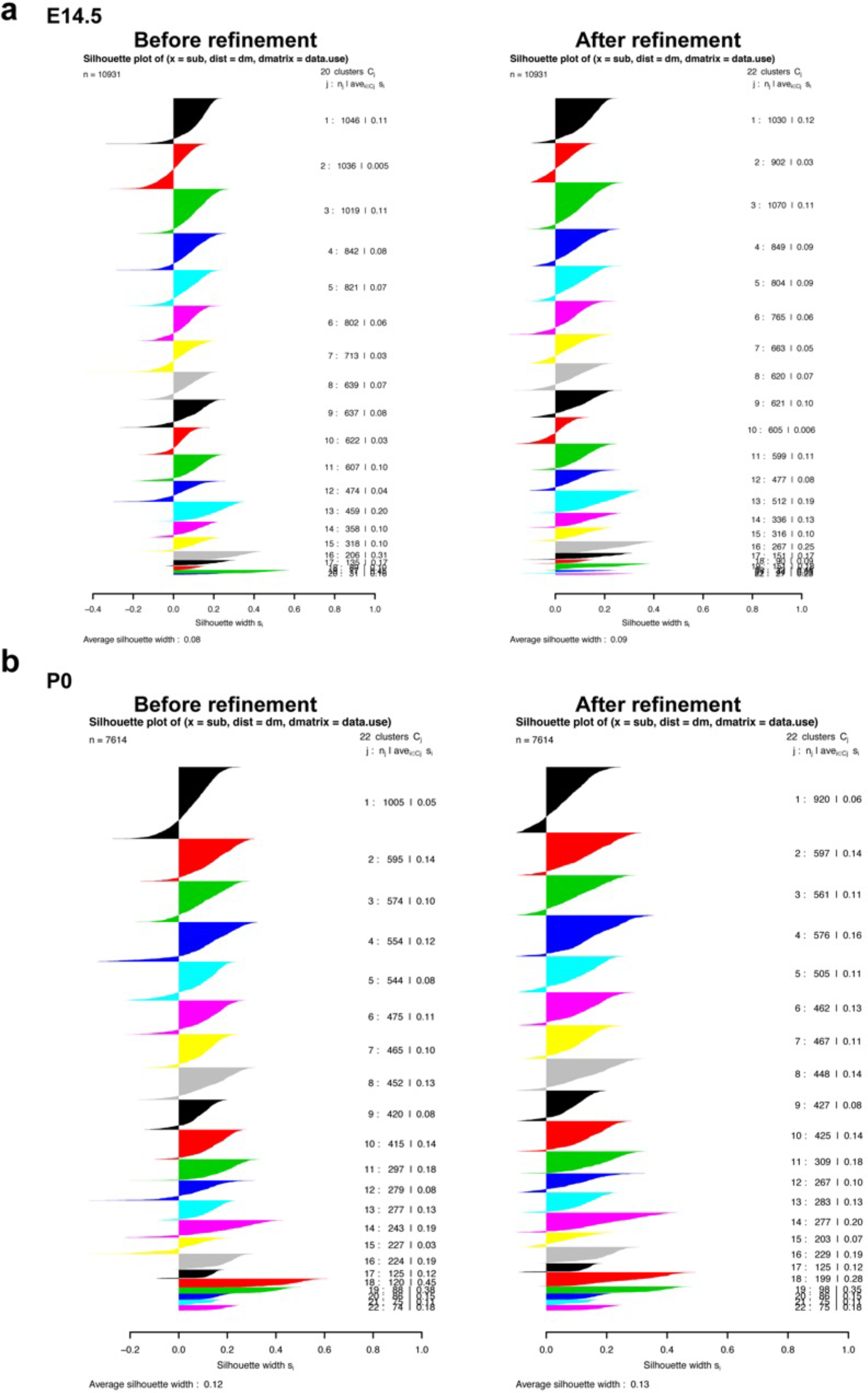
Cluster refinement methodology improves cluster assignments and robustness. Silhouette width plots before (left) and after (right) running the cluster refinement method for E14.5 (**a**) and P0 (**b**) samples. Each row represents one cell, and the silhouette widths were determined using Spearman correlation distances. Positive values indicate a robust cluster assignment. The method allows some cluster outliers to form their own novel clusters, which happens twice in the E14.5 samples.

**Supplementary Figure 4.**
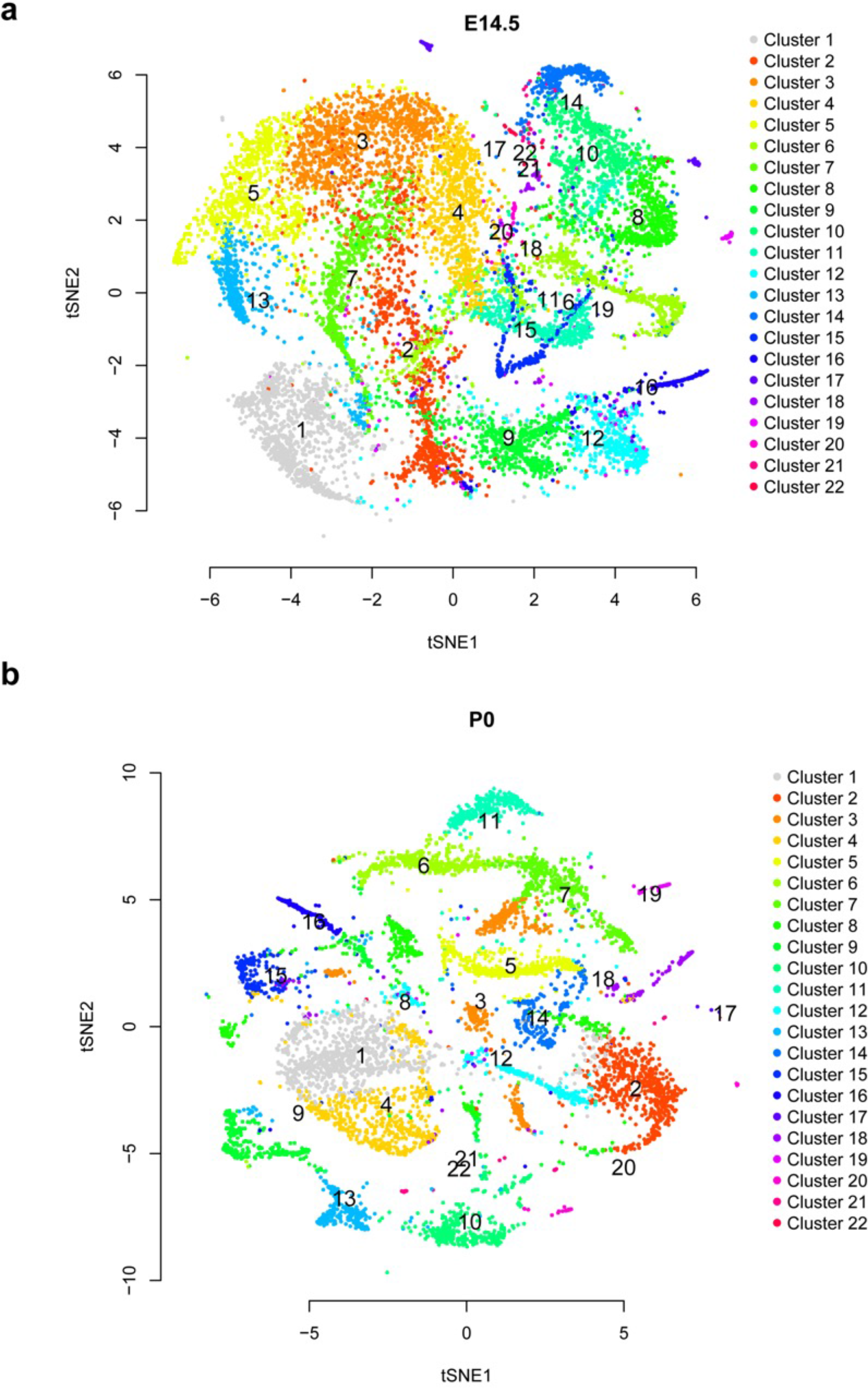
Drop-seq expression data visualized by tSNE. tSNE visualization of significant principal components, where points are labeled by their final cluster assignments for E14.5 (**a**) and P0 (**b**).

**Supplementary Figure 5.**
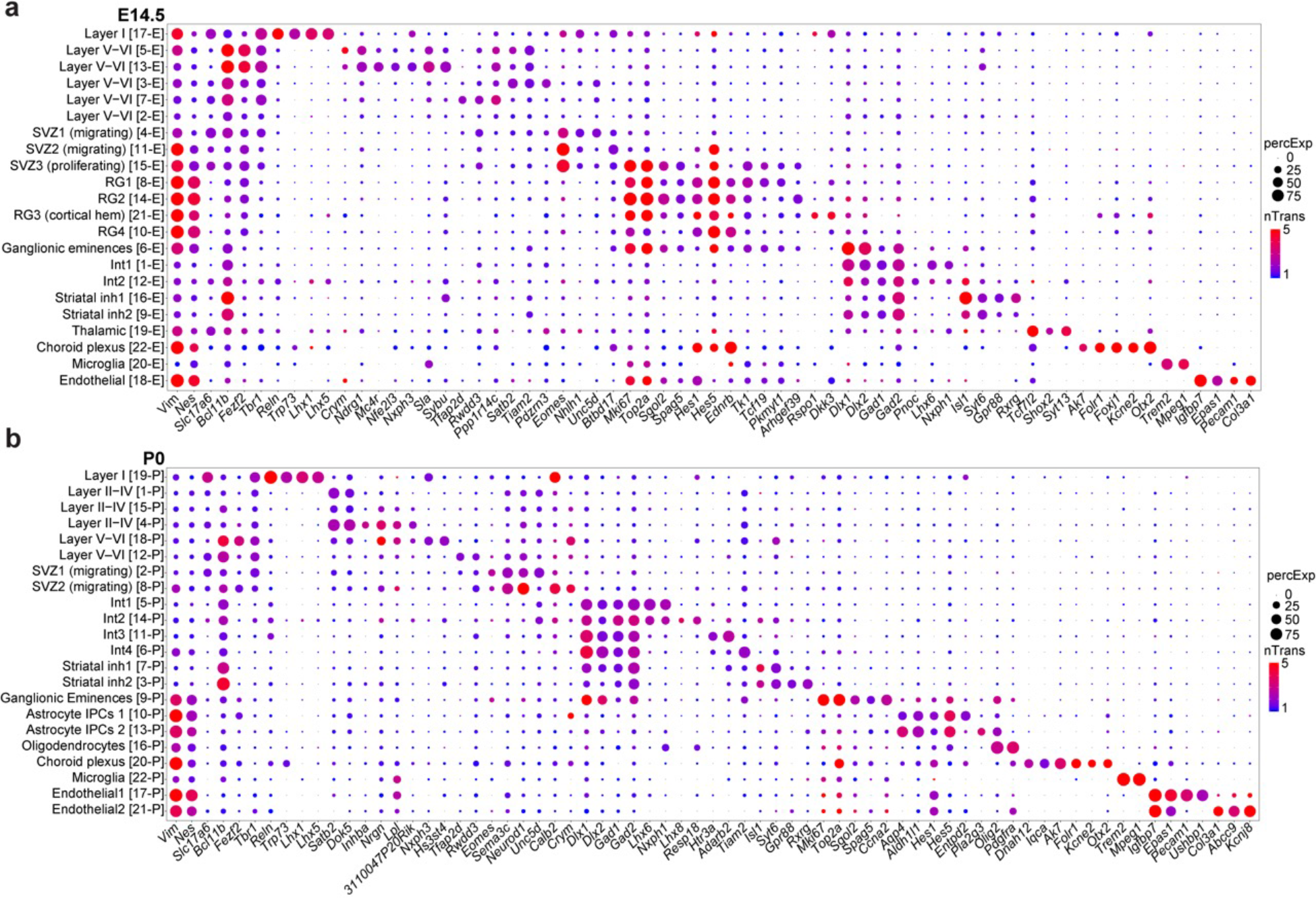
Functional characterization of cell types using expression of known and novel marker genes. Dotplot representations of expression levels and proportion of cells with detected transcripts for markers of each cell type for each age, E14.5 (**a**), and P0 (**b**). Large and red circles indicate expression was high in a large proportion of cells in that cluster, small and blue circles indicate expression was low and detected in few cells in that cluster.

**Supplementary Figure 6.**
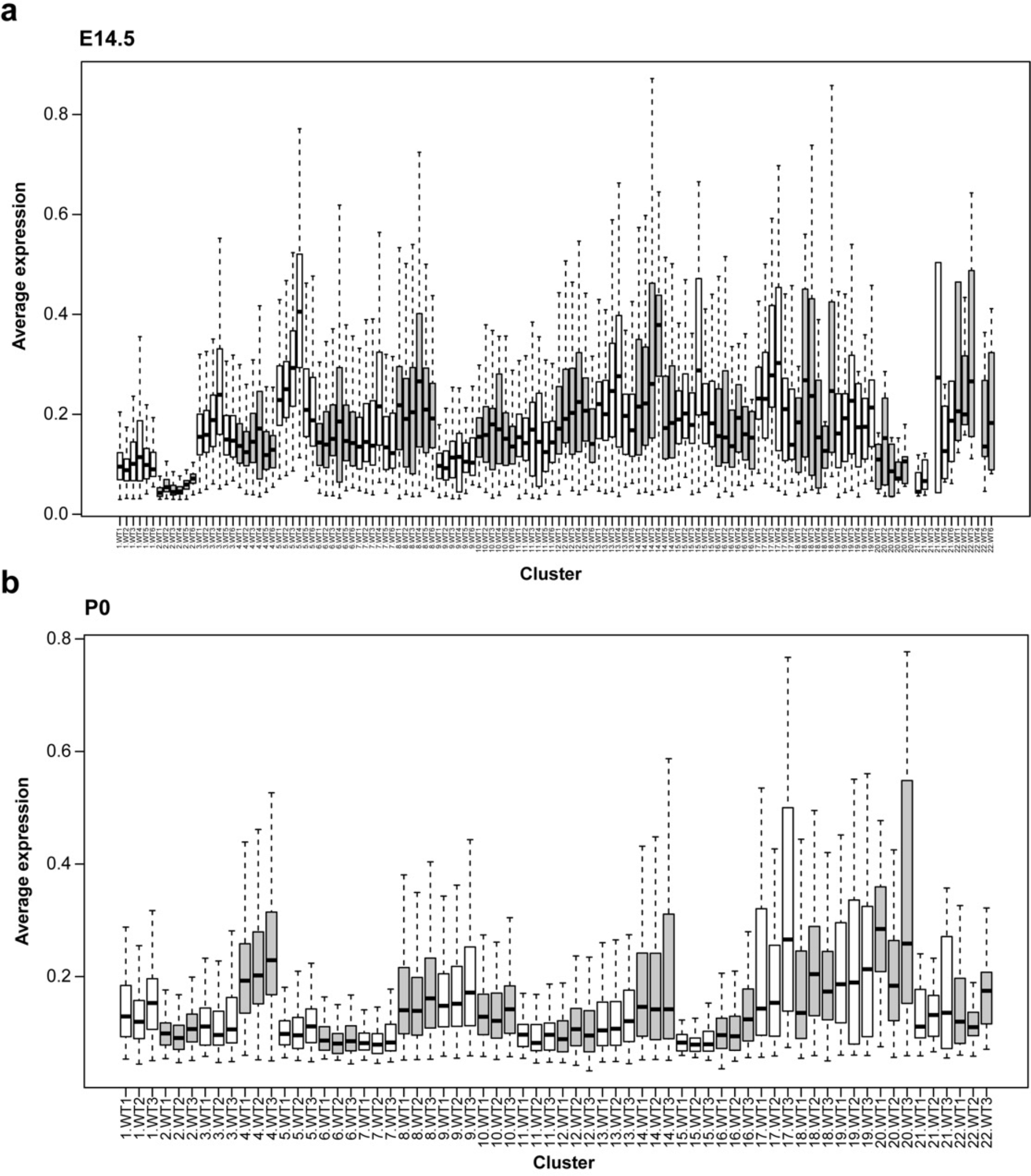
Cross-replicate comparison of library quality, separated by cluster. Box-and-whisker plots depicting the distribution of the average expression values for each biological replicate for each cluster, for E14.5 (**a**), and P0 (**b**). Expression values for all genes for a given biological replicate were averaged within each cluster. Variability cluster-to-cluster is expected, as different cell types may have differing numbers of transcripts expressed, however, variability across replicates within a given cluster is limited, suggestive of consistent representation and strong reproducibility.

**Supplementary Figure 7.**
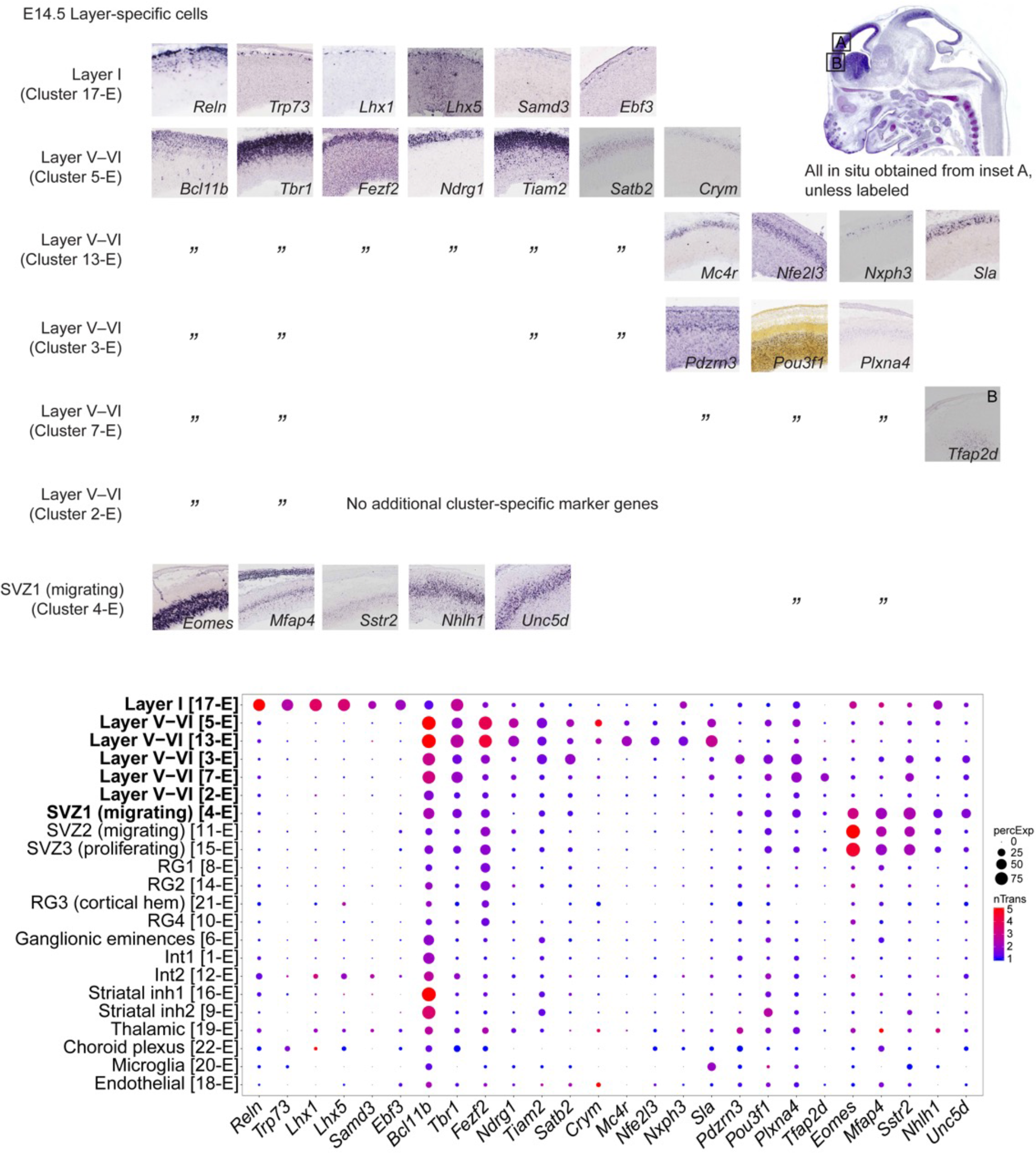
Validation of marker gene expression in E14.5 layer-specific cells. RNA *in situ* hybridization images were obtained from E14.5 cerebral cortex, provided by Eurexpress online database (www.eurexpress.org) except *Pou3f1* obtained from Allen Institute for Brain Science (www.developingmouse.brain-map.org). Layer I (17-E) expresses Cajal-Retzius cell markers *Reln*, *Trp73*, *Lhx1* and *Lhx5*. Layer V-VI clusters (5-E, 13-E, 3-E, 7-E and 2-E) express deep layer markers, *Bcl11b* (also known as *Ctip2*) and *Tbr1*. Expression of *Satb2* in clusters 5-E, 13-E and 3-E suggests eventual differentiation into upper layer callosal projection neurons. SVZ1 (4-E) expresses *Eomes* (also known as *Tbr2*), *Sstr2*, a migrational marker, and *Unc5d*, a netrin receptor involved in cell migration. Dotplot representations of expression levels and proportion of cells with detected transcripts for markers of each cell type. Y axis denotes cluster type; X axis denotes gene name. See Supplementary Table 3 for annotation of selected markers.

**Supplementary Figure 8.**
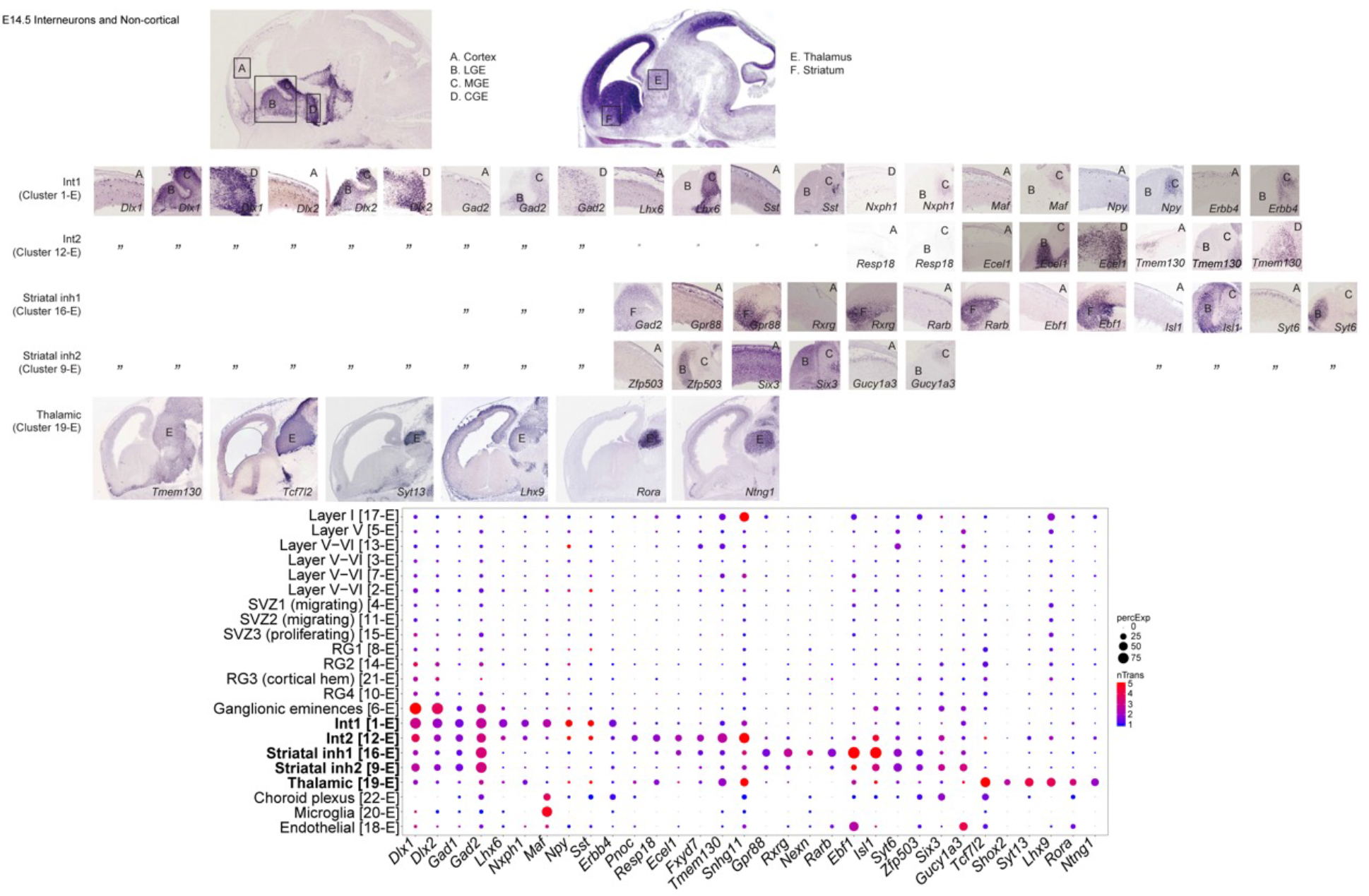
Validation of marker gene expression in E14.5 interneurons and non-cortical cells. RNA *in situ* hybridization images were obtained from E14.5 cerebral cortex, ganglionic eminences, striatum and thalamus, provided by Eurexpress online database (www.eurexpress.org). Int1 (cluster 1-E) expresses *Lhx6*, a transcription factor expressed in MGE-derived cortical Parvalbumin (PV) and Somatostatin (SST) interneurons. Int2 (cluster 12-E) expresses lower levels of Lhx6 and RNA *in situ* expression of *Resp18*, *Ecel1* and *Tmem180* indicate a second distinct population of PV and SST cortical interneurons. Sst is expressed in these 2 clusters but *Pvalb* is not detected. 16-E and 9-E express *Isl1*, a lateral ganglionic eminence (LGE) marker. A subset of these LGE-derived GABAergic neurons are possible cortical interneurons^15^. However, 16-E expresses striatal markers *Gpr88*, *Rxrg* and *Rarb* while 9-E expresses *Zfp503,* a marker of developing striatum and are therefore designated as Striatal inh1 and 2. A small population of cells (1.28% of total population, cluster 19-E) express thalamic markers *Tcf7l2*, *Shox2*, *Syt13*, *Lhx9*, *Rora* and *Ntng1*. Dotplot representations of expression levels and proportion of cells with detected transcripts for markers of each cell type. Y axis denotes cluster type; X axis denotes gene name. See Supplementary Table 3 for annotation of selected markers.

**Supplementary Figure 9.**
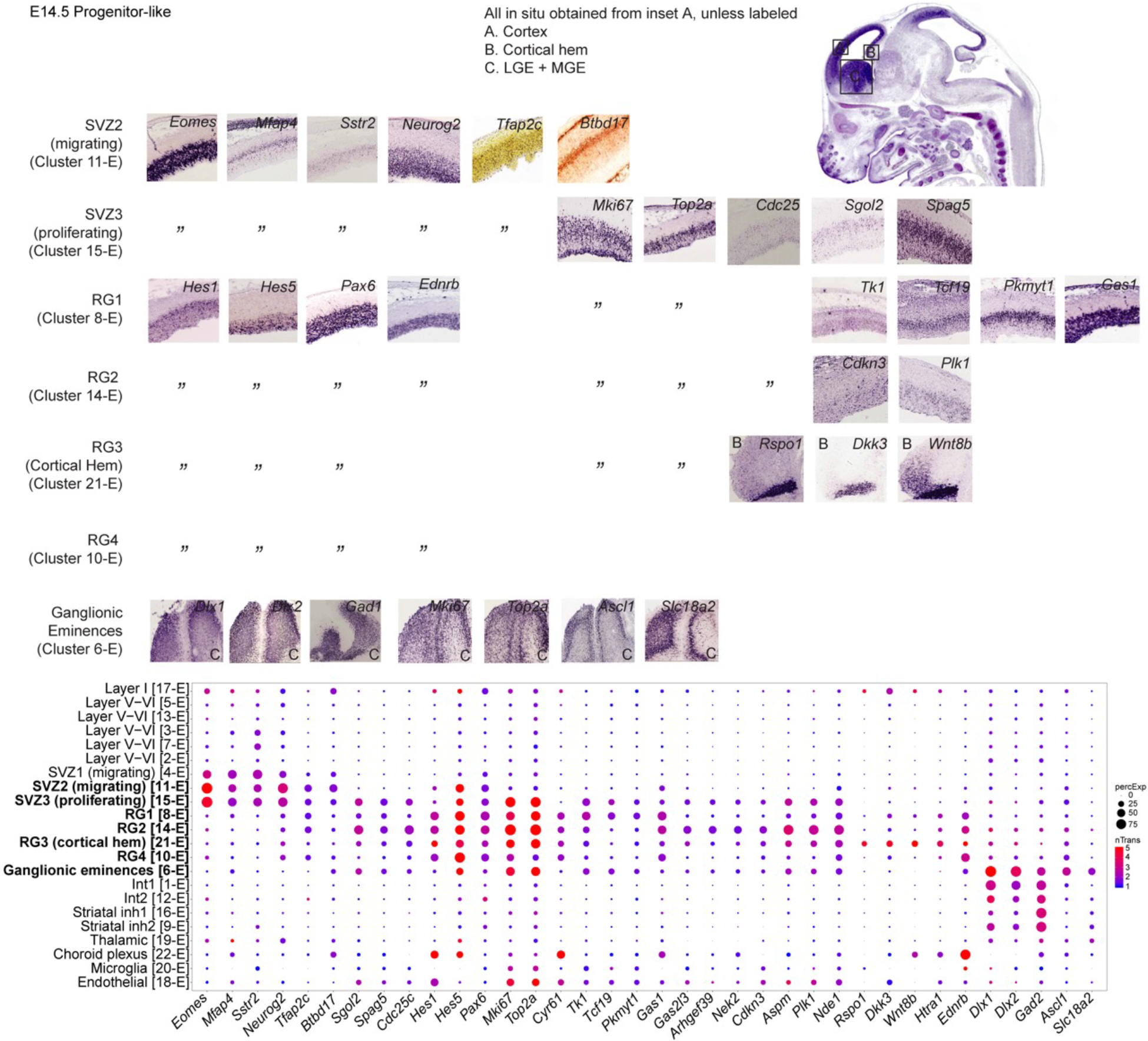
Validation of marker gene expression in E14.5 progenitor-like cells. RNA *in situ* hybridization images were obtained from E14.5 cerebral cortex, cortical hem and ganglionic eminences, provided by Eurexpress online database (www.eurexpress.org) except *Tfap2c* obtained from Allen Institute for Brain Science (www.developingmouse.brain-map.org) and *Btbd17* from GENSAT online database (www.gensat.org). SVZ2 (11-E) expresses *Sstr2*, a migrational marker and *Neurog2*, which initiates delamination of radial glia and neuronal lineage commitment. SVZ3 (15-E), RG1 (8-E), RG2 (14-E), RG3 (21-E) and ganglionic eminences (6-E) express *Mki67* and *Top2a,* markers of cell proliferation. RG1-4 express radial glia markers *Hes1, Hes5* and *Pax6,* while RG3 specifically, expresses *Rspo1*, *Dkk3*, and *Wnt8b*, markers of cortical hem. Ganglionic eminences (6-E) express various GABAergic markers such as *Dlx1*, *Dlx2* and *Gad2*. Dotplot representations of expression levels and proportion of cells with detected transcripts for markers of each cell type. Y axis denotes cluster type; X axis denotes gene name. See Supplementary Table 3 for annotation of selected markers.

**Supplementary Figure 10.**
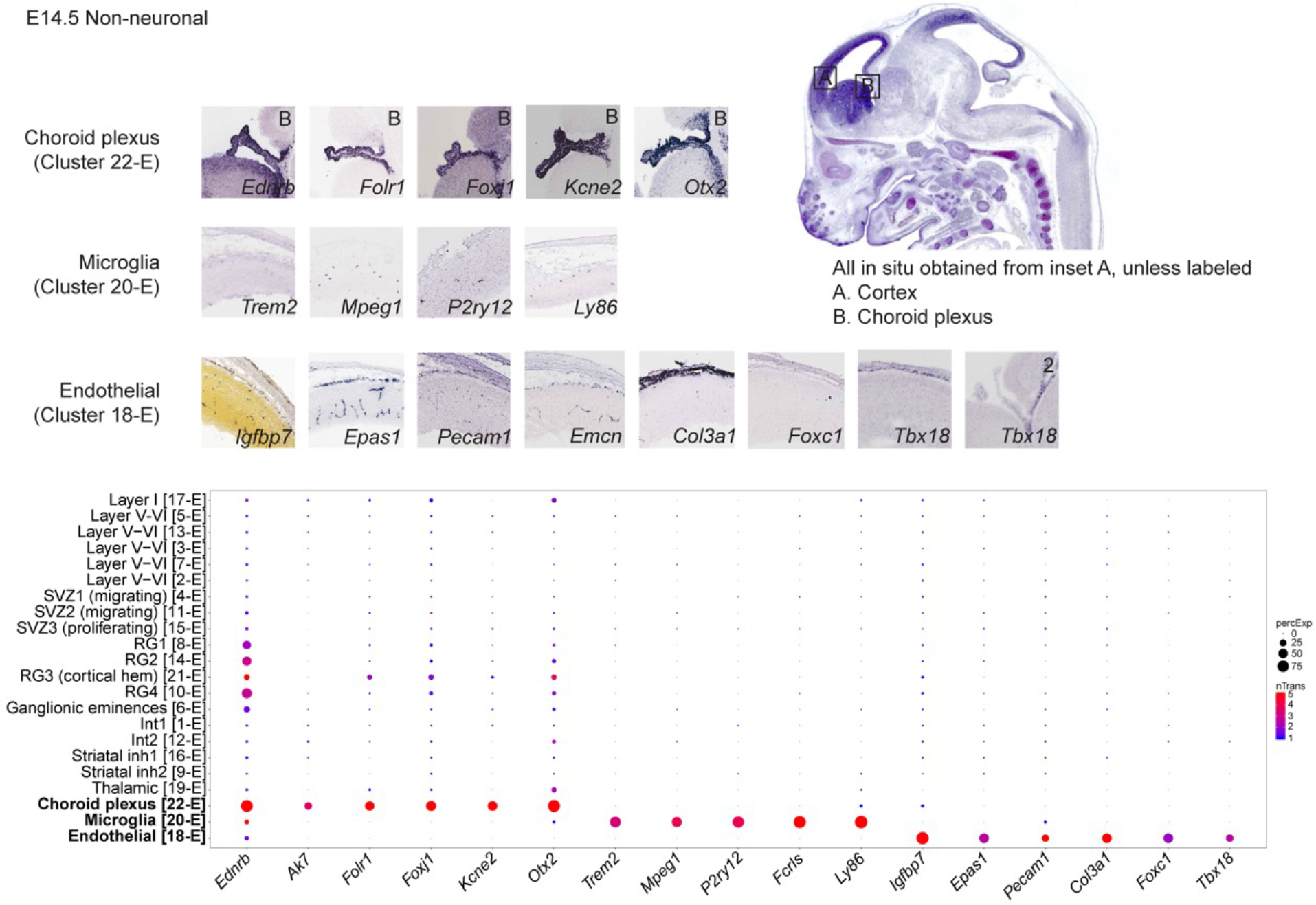
Validation of marker gene expression in E14.5 nonneuronal cells. RNA *in situ* hybridization images were obtained from E14.5 cerebral cortex and choroid plexus, provided by Eurexpress online database (www.eurexpress.org) except *Igfbp7* obtained from Allen Institute for Brain Science (www.developingmouse.brain-map.org). Cluster 22-E expresses choroid plexus markers *Folr1*, *Foxj1*, *Kcne2* and *Otx2*. Cluster 20-E expresses microglial markers *Trem2*, *Mpeg1*, *P2ry12* and *Ly86*. Cluster 18-E expresses endothelial markers *Igfbp7*, *Epas1* and *Pecam1*. Dotplot representations of expression levels and proportion of cells with detected transcripts for markers of each cell type. Y axis denotes cluster type; X axis denotes gene name. See Supplementary Table 3 for annotation of selected markers.

**Supplementary Figure 11.**
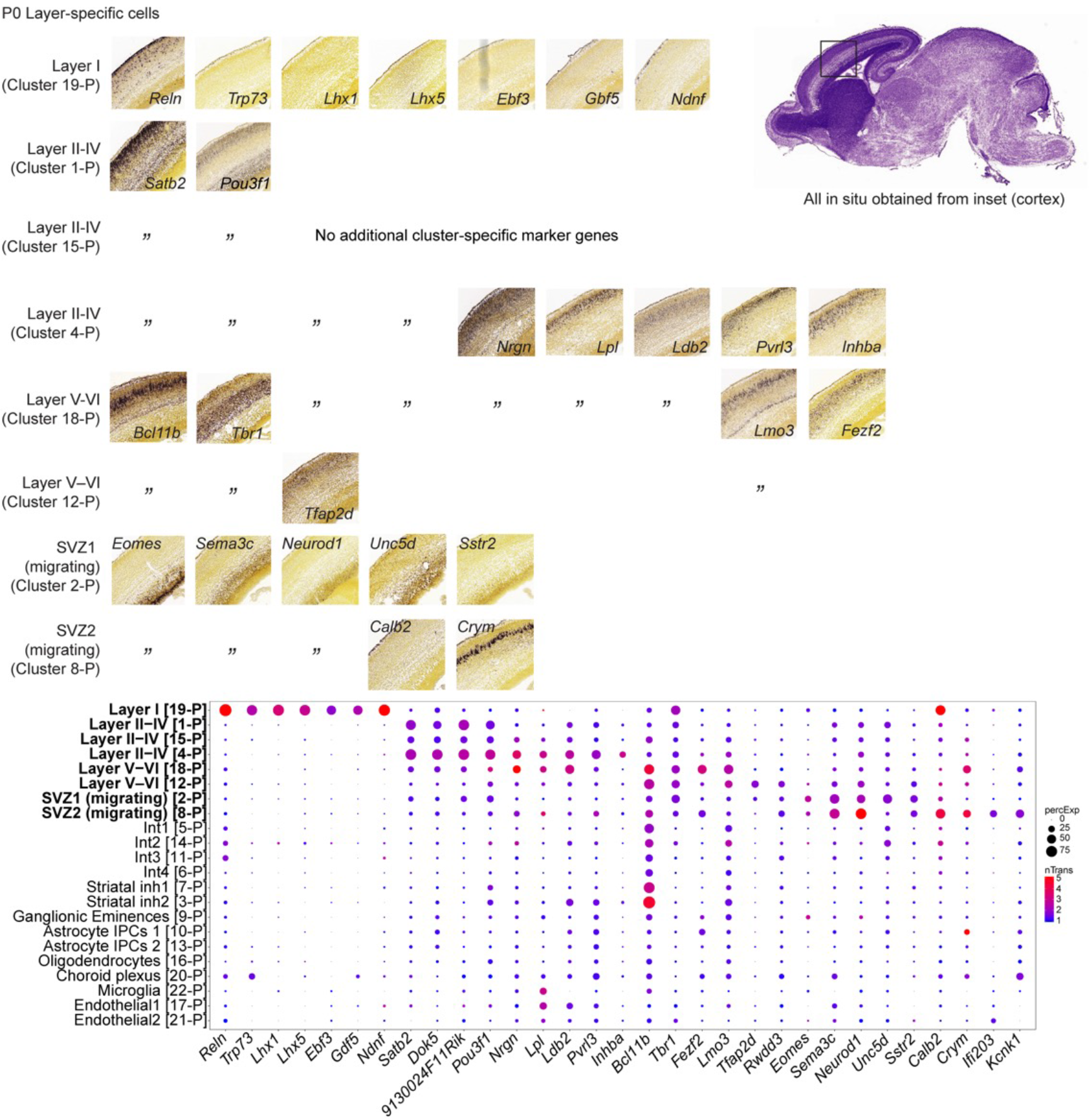
Validation of marker gene expression in P0 layer-specific cells. RNA *in situ* hybridization images were obtained from E18.5 cerebral cortex, provided by Allen Institute for Brain Science (www.developingmouse.brain-map.org). Layer I (cluster 19-P) expresses Cajal-Retzius cell markers *Reln*, *Trp73*, *Lhx1* and *Lhx5*. Layer II-IV (clusters 1-P, 15-P, 4-P) expresses *Satb2*, a marker for upper layer callosal projection neurons and *Pou3f1,* a migrational marker observed in cluster 3-E (Supplementary Figure 7). Deeper layer clusters 18-P and 12-P express *Bcl11b* and *Tbr1*. SVZ1 (2-P) and SVZ2 (8-P) express SVZ marker, *Eomes* (*Tbr2*) and *Sema3c*, a marker of migrating newborn neurons. Dotplot representations of expression levels and proportion of cells with detected transcripts for markers of each cell type. Y axis denotes cluster type; X axis denotes gene name. See Supplementary Table 3 for annotation of selected markers.

**Supplementary Figure 12.**
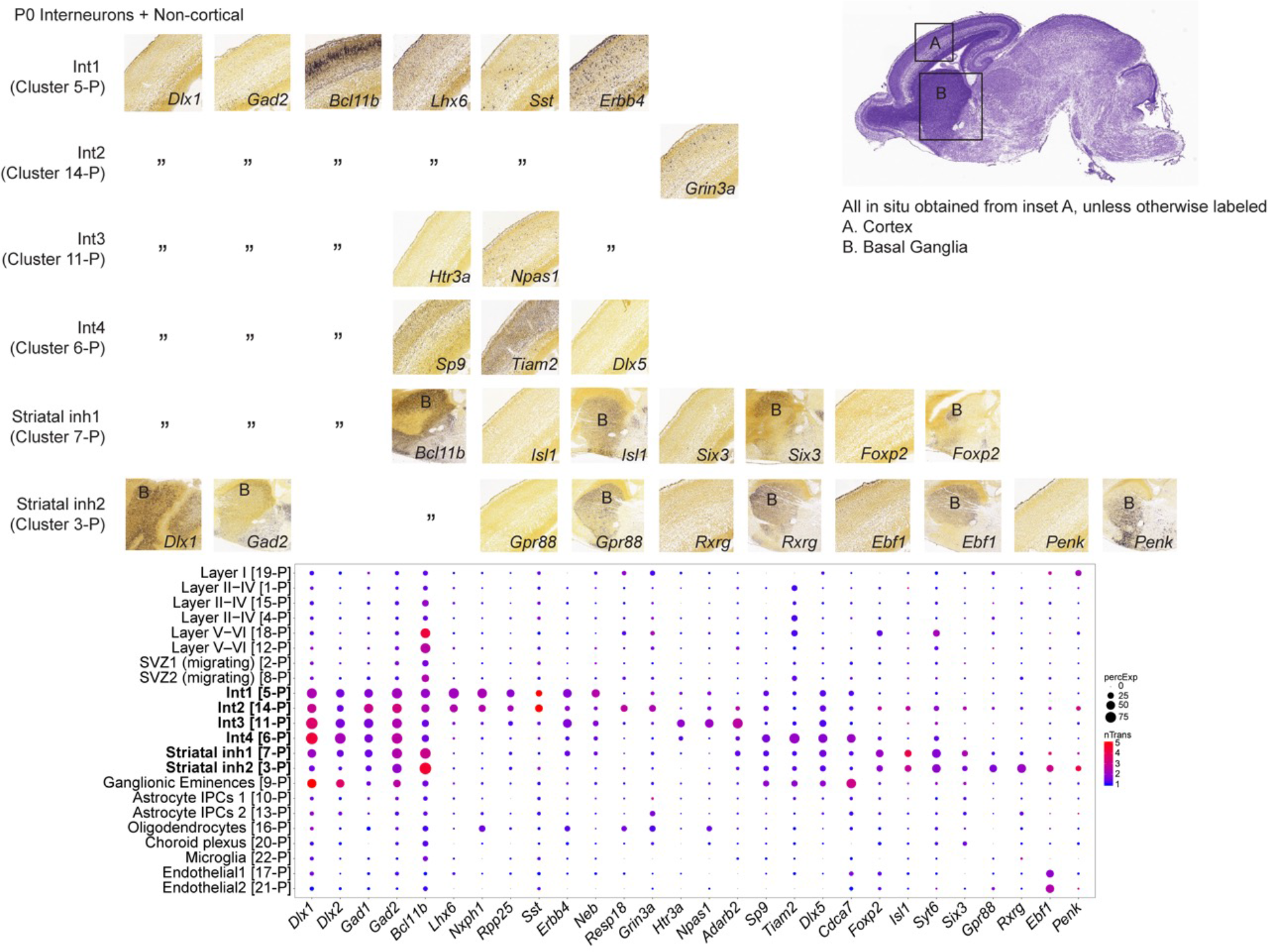
Validation of marker gene expression in P0 interneurons and non-cortical cells. RNA *in situ* hybridization images were obtained from E18.5 cerebral cortex, basal ganglia and preoptic area, provided by Allen Institute for Brain Science (www.developingmouse.brain-map.org). Int1 (5-P), Int2 (14-P), Int3 (11-P), Int4 (6-P), Striatal inh1 (7-P) and Striatal inh2 (3-P) express GABAergic markers *Dlx1*, *Dlx2*, *Gad1* and *Gad2*. Int1 and Int2 express *Lhx6* and *Nxph1*, markers of PV and SST^+^ interneuron precursors, while Int4 expresses *Cdca7*, a marker for some PV and SST^+^ interneurons. Sst is expressed in Int1 and Int2 but *Pvalb* is not detected. Int3 expresses *Htr3a*, *Npas1* and *Adarb2*, markers of VIP cortical interneurons. Striatal inh1 and 2 express striatal markers *Isl1* and *Gpr88* respectively. Dotplot representations of expression levels and proportion of cells with detected transcripts for markers of each cell type. Y axis denotes cluster type; X axis denotes gene name. See Supplementary Table 3 for annotation of selected markers.

**Supplementary Figure 13.**
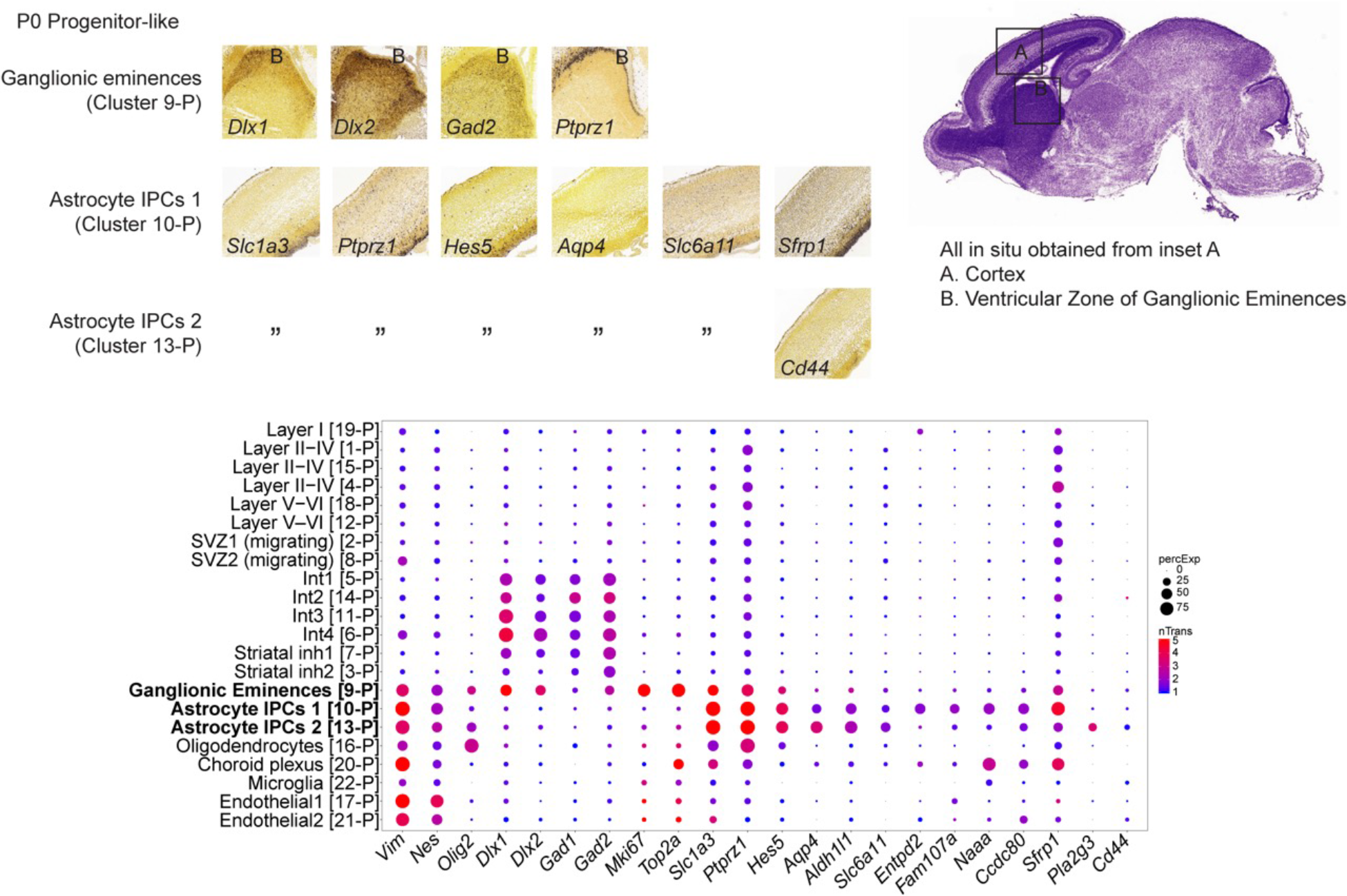
Validation of marker gene in P0 progenitor-like cells. RNA *in situ* hybridization images were obtained from E18.5 cerebral cortex and ganglionic eminence, provided by Allen Institute for Brain Science (www.developingmouse.brain-map.org). Ganglionic eminences (cluster 9-P) express GABAergic markers *Dlx1*, *Dlx2* and *Gad2* and proliferation markers *Mki67* and *Top2a*. Astrocyte IPCs 1 (10-P) and 2 (13-P) express astrocytic markers *Ptprz1, Aqp4* and *Aldh1l1* while maintaining some expression of radial glia marker *Hes5*. Dotplot representations of expression levels and proportion of cells with detected transcripts for markers of each cell type. Y axis denotes cluster type; X axis denotes gene name. See Supplementary Table 3 for annotation of selected markers.

**Supplementary Figure 14.**
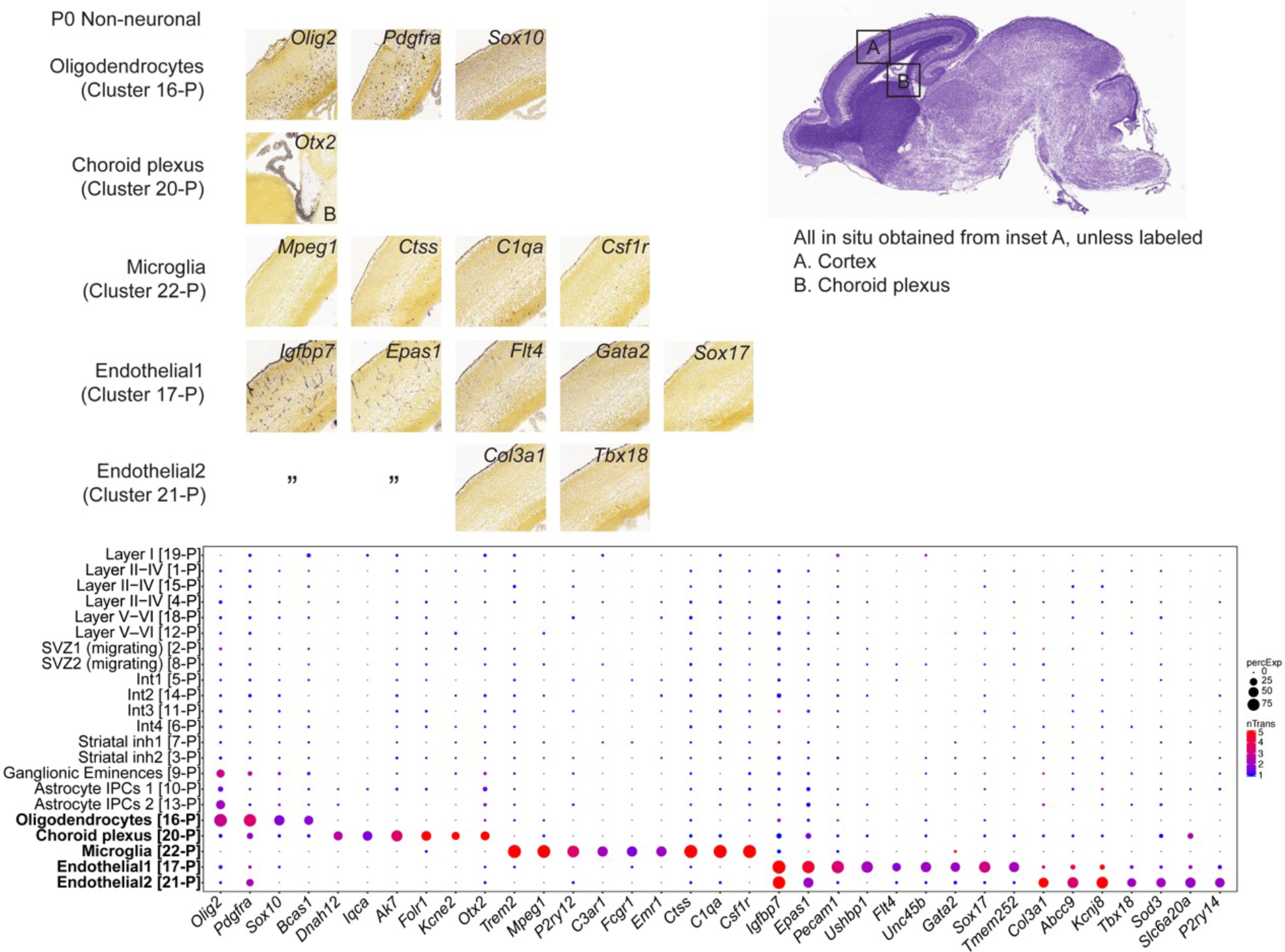
Validation of marker gene expression in P0 nonneuronal cells. RNA *in situ* hybridization images were obtained from E18.5 cerebral cortex and choroid plexus, provided by Allen Institute for Brain Science (www.developingmouse.brain-map.org). Cluster 16-P expresses oligodendrocyte markers *Olig2* and *Pdgfra*. Cluster 20-P expresses choroid plexus markers *Ak7*, *Folr1*, *Kcne2* and *Otx2*. Cluster 22-P expresses microglial markers *Mpeg1*, *Ctss*, *C1qa* and *Csf1r*. Clusters 17-P and 21-P express endothelial markers *Igfbp7* and *Epas1*. Dotplot representations of expression levels and proportion of cells with detected transcripts for markers of each cell type. Y axis denotes cluster type; X axis denotes gene name. See Supplementary Table 3 for annotation of selected markers.

**Supplementary Figure 15.**
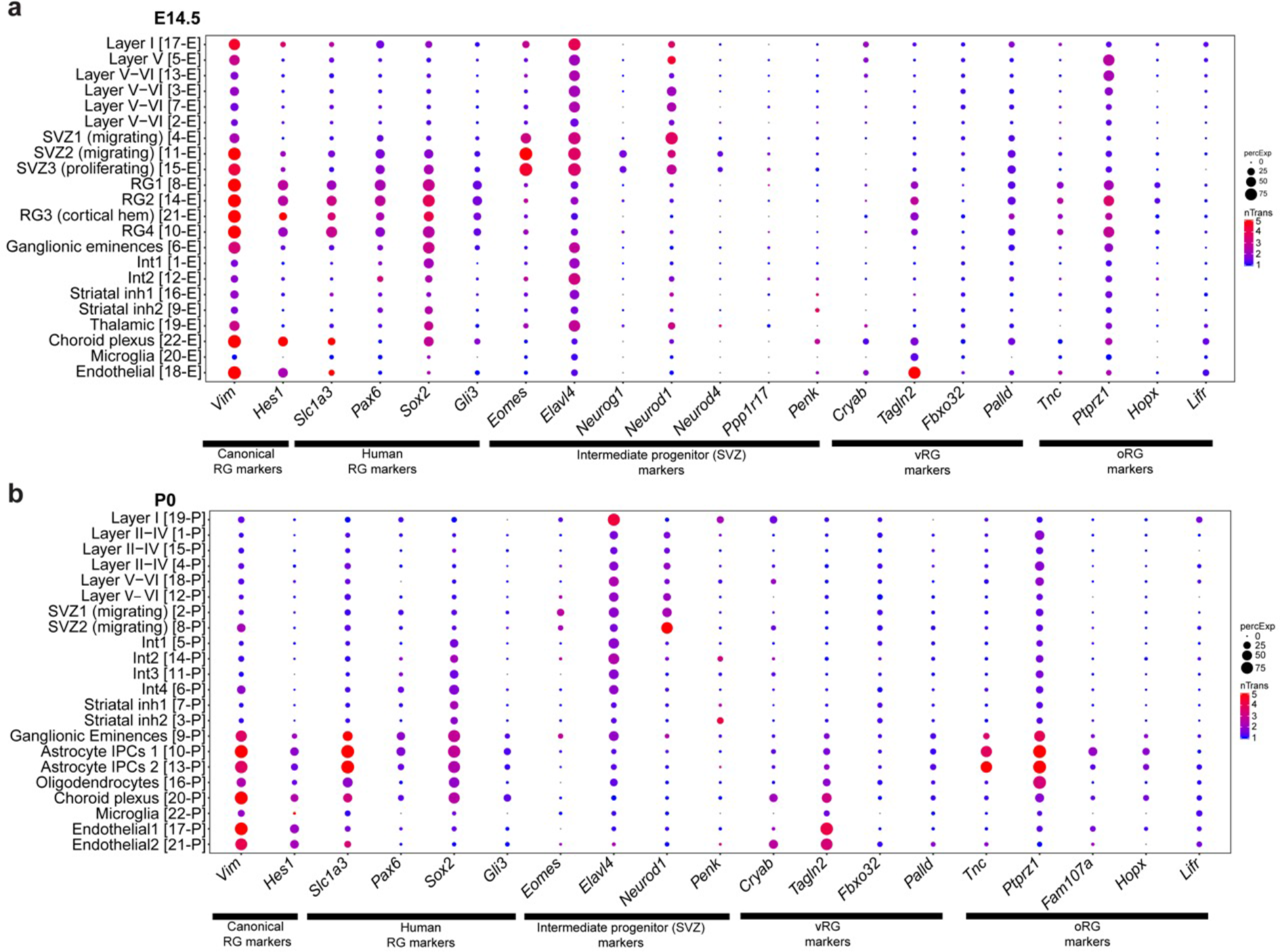
Human radial glia marker gene expression (Pollen et al.)^16^ in developing mouse cortex. Dotplot representations of expression levels and proportion of cells with detected transcripts for markers of canonical radial glia (*Vim*, *Hes1*), human radial glia (*Slc1a3*, *Pax6*, *Sox2*, *Gli1*), intermediate progenitors (*Eomes*, *Elavl4*, *Neurod1*, *Neurod4*, *Ppp1r17*, *Penk*), ventral radial glia (*Cryab*, *Tagln2*, *Fbxo32*, *Palld*) and outer radial glia (*Tnc*, *Ptprz1*, *Fam107a*, *Hopx*, *Lifr*) for each age, E14.5 (**a**), and P0 (**b**). *Neurod4* and *Ppp1r17* expression were not detected at E14.5 while *Fam107a* was not detected at P0. Large and red circles indicate expression was high in a large proportion of cells in that cluster, small and blue circles indicate expression was low and detected in few cells in that cluster.

**Supplementary Figure 16.**
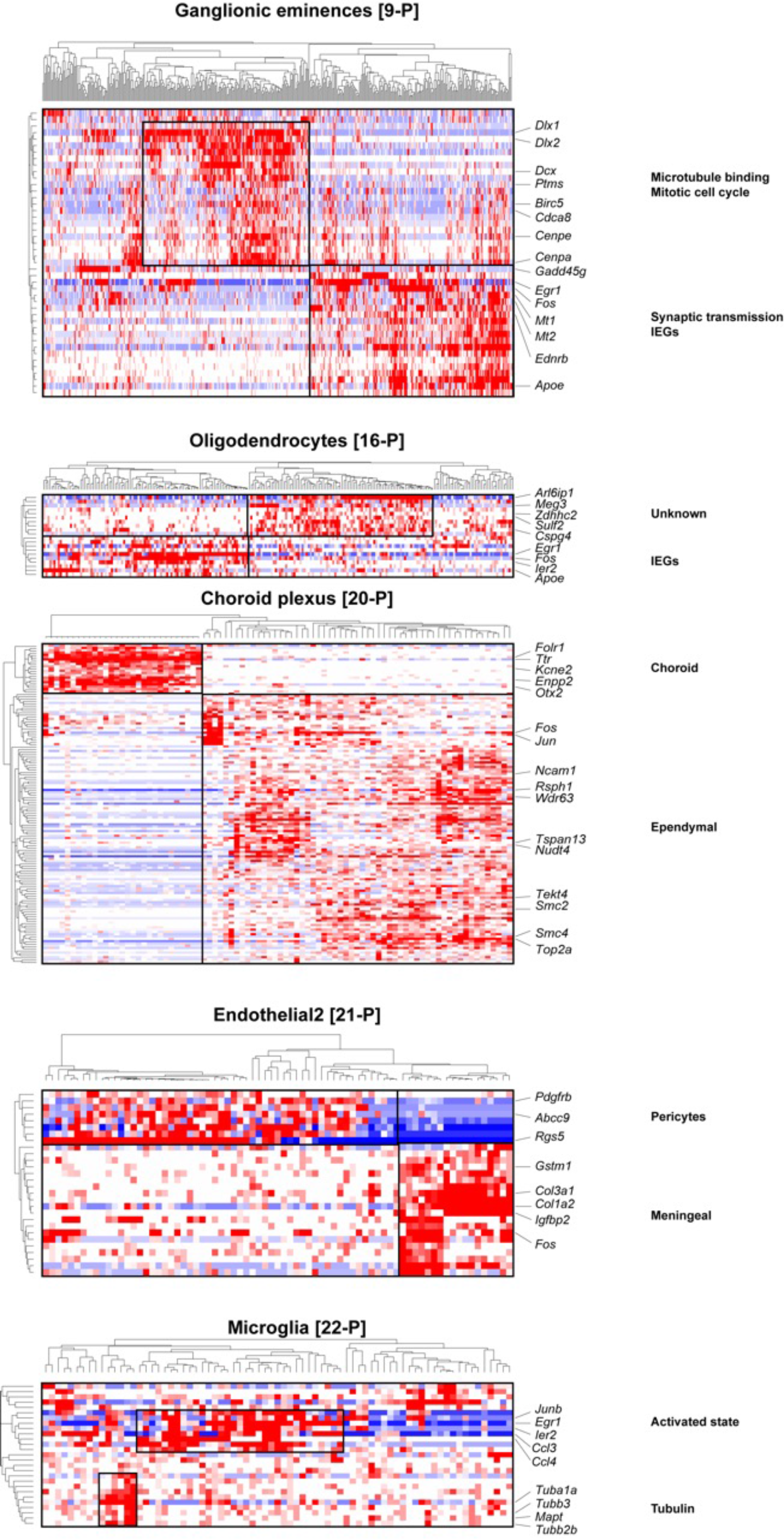
Heatmap representation of cellular sub-clusters within E14.5 cell types. We identified cellular sub-clusters in 7 of the 22 E14.5 cell types. The expression of sub-cluster-specific genes was median-centered and hierarchically clustered for all cells within the greater cluster and expression values were then plotted as a heatmap. Some key genes of interest and functional ontologies/pathways into which these genes fall are shown. Marker genes that describe the entire cluster or other genes expression in >75% or <25% of cells are not shown.

**Supplementary Figure 17.**
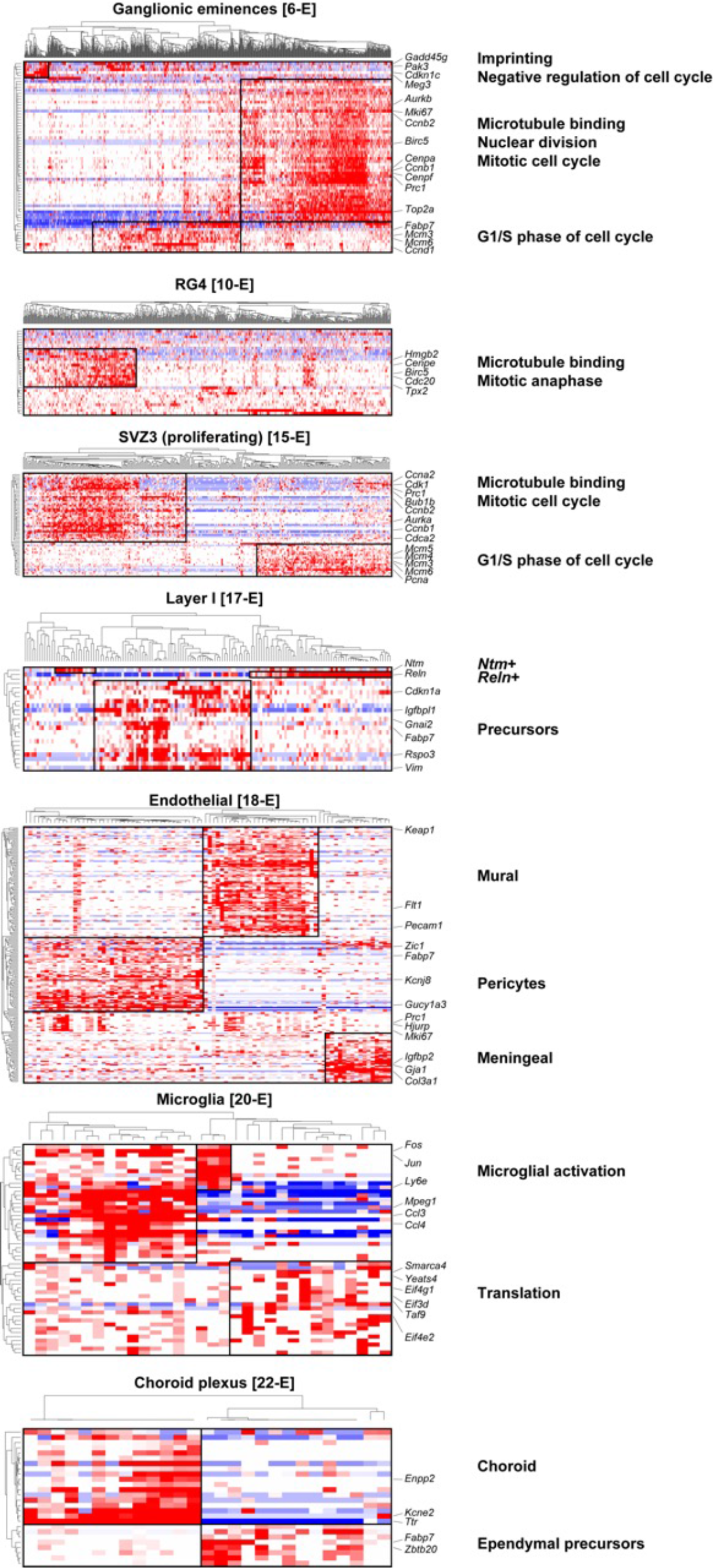
Heatmap representation of cellular sub-clusters within P0 cell types. We identified cellular sub-clusters in 5 of the 22 P0 cell types. The expression of sub-cluster-specific genes was median-centered and hierarchically clustered for all cells within the greater cluster and expression values were then plotted as a heatmap. Some key genes of interest and functional ontologies/pathways into which these genes fall are shown. Marker genes that describe the entire cluster or other genes expression in >75% or <25% of cells are not shown.

**Supplementary Figure 18.**
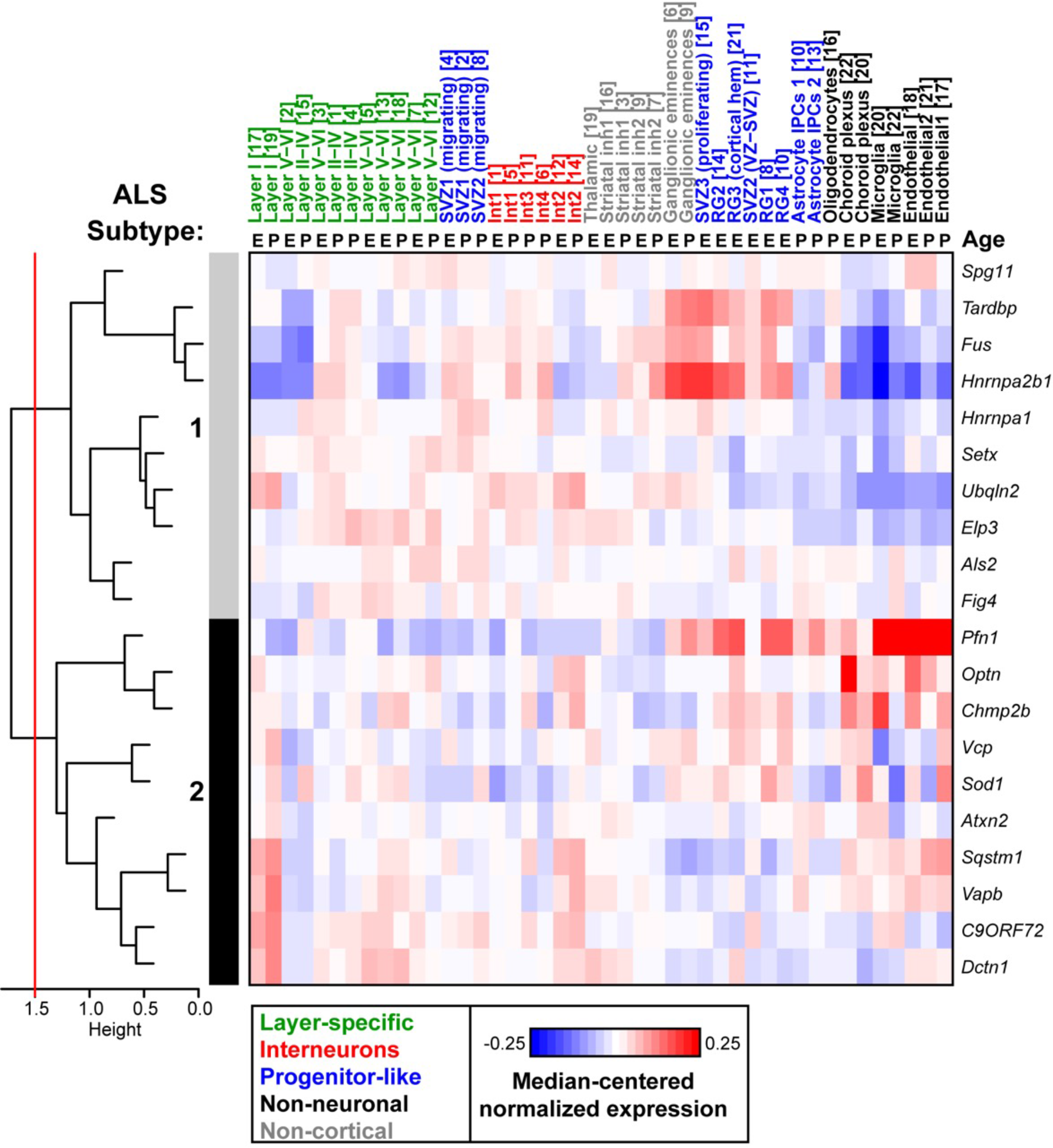
Genes linked to amyotrophic lateral sclerosis (ALS) form two subtypes based on cortical expression patterns. Heatmap of ALS-linked genes clustered hierarchically by their expression across all 22 cell types for both E14.5 and P0. Pearson correlation distances were used to draw the dendrogram, and a height threshold of 1.5 (red line) was used to identify potential subtypes.

**Supplementary Figure 19.**
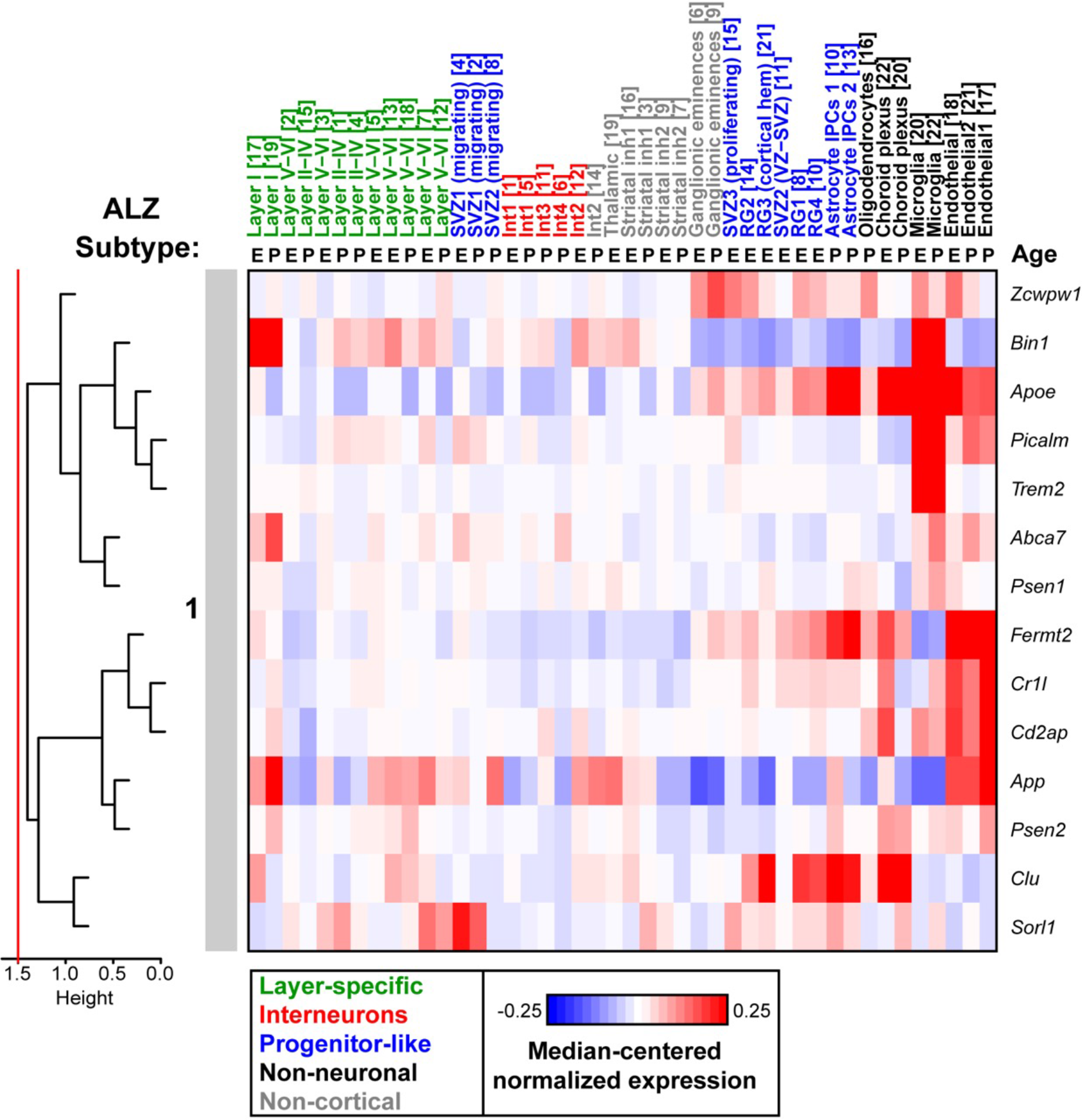
Genes linked to Alzheimer’s disease (ALZ) form one subtype based on cortical expression patterns. Heatmap of Alzheimer’s-linked genes clustered hierarchically by their expression across all 22 cell types for both E14.5 and P0. Pearson correlation distances were used to draw the dendrogram, and a height threshold of 1.5 (red line) was used to identify potential subtypes.

**Supplementary Figure 20.**
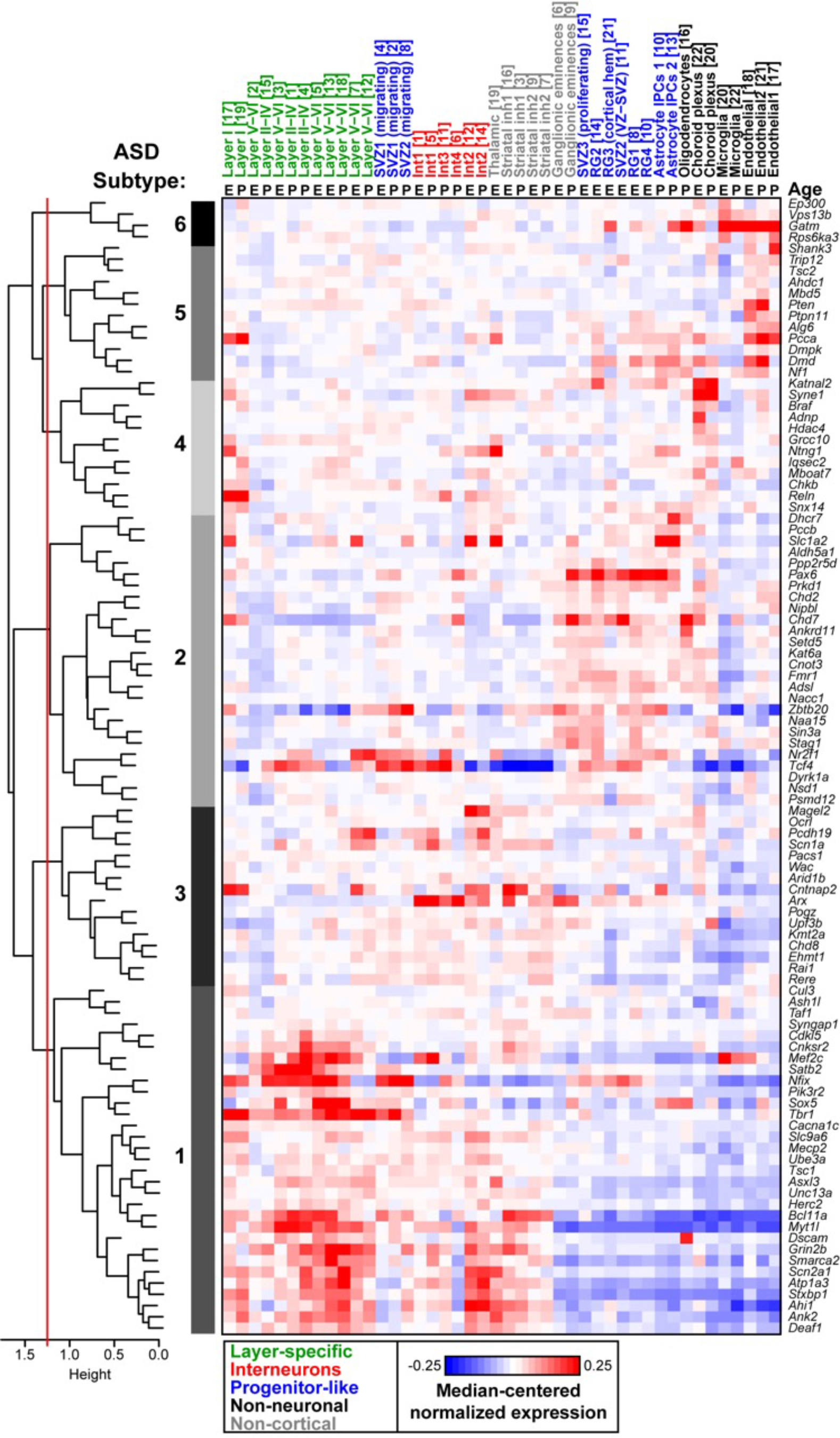
Genes linked to autism spectrum disorder (ASD) form six subtypes based on cortical expression patterns. Heatmap of ASD-linked genes clustered hierarchically by their expression across all 22 cell types for both E14.5 and P0. Pearson correlation distances were used to draw the dendrogram, and a height threshold of 1.25 (red line) was used to identify potential subtypes.

**Supplementary Figure 21.**
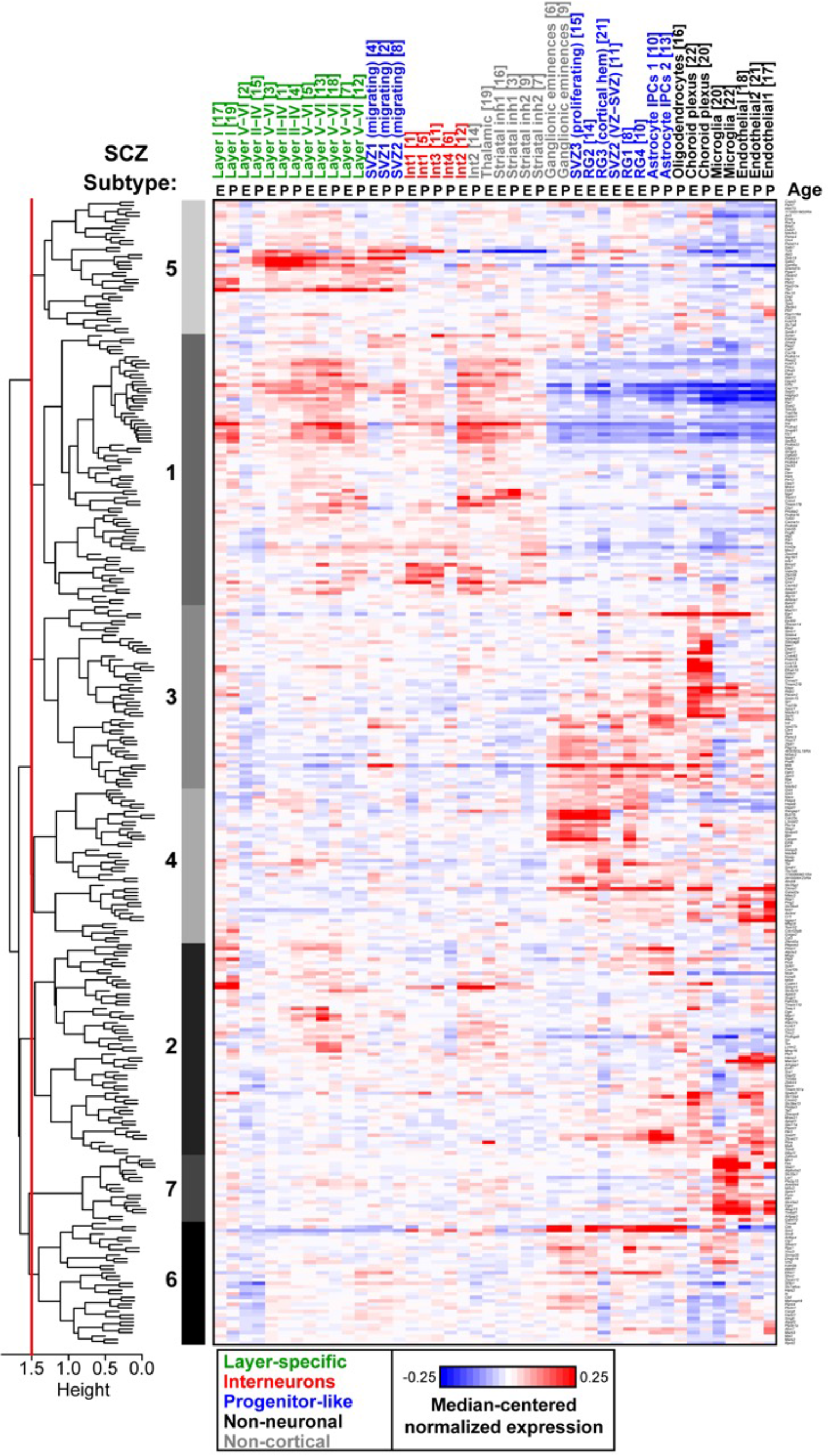
Genes linked to schizophrenia (SCZ) form seven subtypes based on cortical expression patterns. Heatmap of SCZ-linked genes clustered hierarchically by their expression across all 22 cell types for both E14.5 and P0. Pearson correlation distances were used to draw the dendrogram, and a height threshold of 1.5 (red line) was used to identify potential subtypes.

